# Incongruent phylogenies and its implications for the study of diversification, taxonomy and genome size evolution of *Rhododendron* (Ericaceae)

**DOI:** 10.1101/2020.07.27.216218

**Authors:** Gulzar Khan, Jennifer Nolzen, Hartwig Schepker, Dirk C. Albach

## Abstract

**PREMISE:** *Rhododendron* L. (Ericaceae Juss.), is the most species-rich genus of woody flowering plants with > 1000 species. Despite the interest in the genus and numerous previous phylogenetic analysis, the infrageneric classification for the genus is still debated, partly due to its huge diversity, partly due to homoplasy in key characters and partly due to incongruence between phylogenetic markers. Here, we provide a broad coverage of representative species of all *Rhododendron* subgenera, sections, and most subsections to resolve its infrageneric phylogeny or highlight areas of incongruence, support previous analyses of diversification patterns and establish a relationship between genome size evolution and its diversification.

**METHODS:** We generated sequences of two plastid (*trn*K and *trn*L-F) and two nuclear (ITS and *rpb*2-i) markers for a total of 259 *Rhododendron* species, and used likelihood and Bayesian statistics to analyze the data. We analyzed the markers separately to discuss and understand incongruence among the data sets and among previous studies.

**RESULTS:** We found that the larger a subgenus, the more strongly it is supported as monophyletic. However, the smaller subgenera pose several problems, e.g., *R*. subgen. *Azaleastrum* consists of two sections inferred to be polyphyletic. The main shift to higher diversification in the genus occurred in the Himalayan/SE Asian clade of *R*. subgen. *Hymenanthes*. We found that polyploidy occurs in almost all subgenera but most polyploid species are within *R*. subgen. *Rhododendron* sections *Rhododendron* and *Schistanthe*.

**CONCLUSION:** Whereas previous reports stated that genome sizes of tropical plants are lower than those of colder and temperate regions in angiosperms in general, our study provides evidence for such a shift to small genome-tropical species within a genus. Taken together, we see the merit in the recognition of the five major clades at the sub generic level but given the amount of incongruence a large amount of species cannot be confidently assigned to one of these five clades. Further, genome-wide data will be necessary to assess whether these currently unassignable taxa are independent taxa, assignable to one of the five major clades or whether they are inter-subgeneric hybrids.

---

*Rhododendron* L. (Ericaceae Juss.) is the most species-rich genus of woody flowering plants, placed among the twenty largest plant genera >1000 species (Frodin, 2004). The genus has a worldwide distribution (except limited distribution in Central & South America and Africa) with a center of diversity in China (GBIF; Fang and Ming, 1995; Wu and Raven, 2005; Brown et al., 2006). However, many of these species have a limited distribution and are, thus, threatened. According to Gibbs et al. (2011), about 70% of the species of *Rhododendron* are classified as vulnerable, threatened, endangered or critically endangered. Many of the species in the genus are used as ornamentals and/or used as medicinal plants, with abundant literature on their ethno-medicinal use as anti-inflammatory agents, pain killers, in gastro-intestinal disorders, common cold, asthma, skin diseases, and toxic agents (used in the form of insecticide or poison; see Popescu and Kopp, 2013 for a review). Innocenti et al. (2010) suggested that some species of *Rhododendron* have antibacterial activities. Rezk et al. (2015) extended this work and demonstrated that there is a higher susceptibility of Gram-positive bacteria (e.g., *Bacillus subtilis* Ehrenberg) and lesser susceptibility of Gram-negative bacteria (e.g., *Escherichia coli* (Migula) Castellani and Chalmers) towards *Rhododendron* leaf extracts.

This interest in the genus as ornamental and medicinal plant has led to a long list of studies investigating its diversification and classification. Linnaeus (1753) classified the species in two genera, *Azalea* (species with five stamens) and *Rhododendron* (species with ten stamens). Over the next century, as the number and diversity of known species increased, additional genera such as *Rhodora* L. (Linnaeus, 1762), *Vireya* Blume (Blume, 1826) and *Anthodendron* Rchb. (Reichenbach, 1827) were proposed. Especially, in the second half of the nineteenth century *Rhododendron* enjoyed an increase of species number based on the botanical exploration of the Himalayas and China (e.g., Hooker, 1849) and starting a rhododendronmania in Europe (Musgrave et al., 1998). Sleumer (1949, 1966) proposed a detailed classification for the genus that proved highly influential until today. He classified all known *Rhododendron* species at that time into five subgenera and 13 sections. Chamberlain et al. (1996), based on a number of more narrowly focused morphological taxonomic studies, refined previous classifications for the genus grouping the species into eight subgenera and 12 sections (Sleumer, 1966; Cullen, 1980; Chamberlain, 1982; Philipson and Philipson, 1986; Judd and Kron, 1995; Chamberlain et al., 1996). One notable difference between both classifications is the inclusion of *R*. sect. *Therorhodion* Maxim. In Chamberlain et al. (1996) as a subgenus of *Rhododendron*, whereas Sleumer (1966), following Small (1914) considered it separate from *Rhododendron*. The classification of Chamberlain et al. (1996) is still widely used by specialists and gardeners (Cox and Cox 1997; Goetsch et al., 2005). Phylogenetic analyses started to have an influence on the classification of *Rhododendron* in the last 30 years. The most important changes are the inclusion of three genera, *Ledum* (Kron and Judd, 1990), *Diplarche* Hook.f. & Thomson and *Menziesia* Sm. (Craven, 2011). With respect to the intrageneric classification, Goetsch et al. (2005), based on DNA-based phylogenetic analyses, suggested a reduction to five subgenera, *R*. subgen. *Therorodion, R*. subgen. *Rhododendron, R*. subgen. *Hymenanthes* (including *R*. subgen. *Pentanthera* sect. *Pentanthera*), *R*. subgen. *Choniastrum* (=*R*. subgen. *Azaleastrum* sect. *Choniastrum*) and *R*. subgen. *Azaleastrum* (incl. *R*. subgen. *Pentanthera* sect. *Sciadorhodion, R*. subgen *Mumeazalea, R*. subgen. *Candidastrum, R*. subgen. *Tsutsusi*; see details in Table S1).

Establishing relationships in *Rhododendron* based on morphological characteristics but also molecular markers has been difficult because of frequent convergence and hybridization between species. Hybridization is considered to have played an important role in the evolution and speciation of *Rhododendron* through homoploid or allopolyploid speciation (Milne et al., 1999 and 2003; Milne and Abbott, 2008). This is reflected clearly by the large number of horticultural hybrids in *Rhododendron* (over 28,000; Leslie, 2004) as well as the occurrence of natural hybridization indicating weak reproductive barriers (Kron et al., 1993; Milne and Abbott, 2008; Milne et al., 1999 and 2003; Zha et al., 2008 and 2010; Zhang et al., 2007). Besides homoploid hybridization, polyploidy, the occurrence of three or more sets of homologous chromosomes in the genome also occurs naturally in *Rhododendron*, ranging from triploids to dodecaploids (Ammal, 1950) and even more extensively exploited in horticulture (Jones et al., 2007). In *Rhododendron* the basic chromosome number is 13 (with exception of *R*. *camtschaticum* Pall. with n = 12) with more than 70% of the species counted (∼15% total) being diploid (2n = 26; Ammal et al., 1950; Väinölä, 2000; and Rice et al., 2015), but little is known about the importance of polyploidization in the diversification of the genus. Though hybridization can play diverse roles in promoting speciation (Abbott et al., 2013; and Milne et al., 2010), in *Rhododendron*, hybrid populations have often been found to show higher fitness than their parents only in mosaic habitats created along altitudinal, radiation (Milne et al., 2003) or soil pH (Milne and Abbott, 2008) gradients.

The large number of species has stirred interest in investigating the underlying reasons for its diversification. Milne et al. (2010) demonstrated that *R*. subgen. *Hymenanthes* (Blume) K. Koch within South East (SE) Asia has been the clade to diversify fastest. Similarly, Schwery et al. (2015) supported that the greatest diversity within *Rhododendron* occurs in the Himalayas and Malesia, detected a nested Himalayan *Rhododendron* radiation of species of *R*. subgen. *Hymenanthes*, and a separate diversification of *R*. section *Schistanthe* Schltr. (= *Vireya*) accompanied by an eastwards dispersal, as predicted by (Brown et al., 2006) and Goetsch et al. (2011). However, hybridization or polyploidy have not been considered by these studies are, until now, insufficiently considered in the analyses trying to understand the diversification of the genus, in contrast to biogeographic processes (Shreshta et al., 2018). A phylogenetic study analyzing the importance of hybridization and establishing a robust infrageneric classification requires as many species as possible representing all subgenera, sections, and subsections. Till to date about 400 species of the more than 1000 *Rhododendron* species known to data have been studied in different phylogenetic studies. However, these studies mostly focused on some specific subgenera and/or sections. Only few studies considered the phylogeny of the whole genus *Rhododendron* and mostly using a single DNA region, e.g., plastid *mat*K - 51 species (Kurashige et al., 2001); nuclear ITS - 21 species (Gao et al., 2002); nuclear *rpb*2 - 88 species (Goetsch et al., 2005) or *trn*K, *trn*L-F & ITS – 87 species (Grimbs et al., 2017). Shrestha et al. (2018) investigated the global distribution and molecular phylogeny of *Rhododendron* in a biogeographical context, using 423 species with a concatenated dataset of nine plastid genes, nuclear ribosomal ITS data and six introns of one nuclear gene (RPB). However, neither did they report how much missing data was included nor did they discuss whether and where incongruence between different data sets occurred. In addition, Shrestha et al. (2018) depicted relationships without support values, which does not allow evaluation of the robustness of relationships. Differences between these phylogenies and previous classifications have raised doubts regarding the validity of the morphology-based *Rhododendron* classification and prompted Goetsch et al. (2005) to propose a new classification. However, the differences between analyses based on different DNA regions at the subgeneric and sectional level prevented widespread support for this alternative classification for *Rhododendron*. To this end, here our objective is to reconstruct the phylogeny of *Rhododendron* using representative species of all subgenera, sections, and subsections for discussing support for alternative classifications, especially the DNA-based classification of Goetsch et al. (2005). We used both plastid and nuclear markers for phylogeny reconstruction. Based on this phylogeny, we analyzed the importance of hybridization and polyploidization in diversification of *Rhododendron* in a phylogenetic context. We provide here a critical evaluation of phylogenetic studies and results from different markers in the genus. We, further, use the results to infer patterns of diversification in the genus focusing on the importance of polyploidy and genome size evolution. Our main questions were: (1) Does expansion of the sampling resolve the phylogenetic relationships within *Rhododendron* and support the monophyly of major clades; (2) Do different DNA markers similarly show and support previous diversification patterns? And (3): Are polyploidy and genome size evolution related to the diversification of the genus? To follow these questions, we generated sequences of two plastid (*trn*K and *trn*L-F) and two nuclear (ITS and *rpb*2-i) markers for a total of 307 individuals from 259 *Rhododendron* species. We analyzed all four datasets separately and discuss the incongruences among these markers and previous studies. In addition, we used BAMM (Bayesian Analyses of Macroevolutionary Mixtures; Rabosky, 2014) to investigate the pattern of diversification in *Rhododendron*.

## MATERIAL AND METHODS

### Sampling

We based our sampling strategy on two criteria, first to include as many species as possible and second to represent all subgenera, sections, and subsections of *Rhododendron* following Chamberlain et al. 1996 (Table S1), which we managed with the exception of two subsections of *R*. subgen. *Hymenanthes* (subsectt. *Barbata, Lanata*), the monotypic *R*. subgen. *Pentanthera* subsect. *Sinensia*, and four monotypic subsections in *R*. subgen. *Rhododendron* (subsect. *Afghanica, Campylopogon, Camelliflorum, Virgata*). We collected fresh leaves of 307 individuals from 259 species at Rhododendron-Park Bremen and downloaded the available sequences of other species from GenBank (Table S2). To root the phylogeny of *Rhododendron*, we used *Calluna vulgaris* (L.) Hull, *Empetrum nigrum* L., *Kalmia angustifolia* L., *Kalmia procumbens* (L.) Desvaux, *Vaccinium* x *intermedium* Ruthe, and *Vaccinium myrtilloides* Michx. as outgroups. The outgroup species belong to the same family, covering the main lineages of family Ericaceae and cultivated in the Botanical Garden of the Carl von Ossietzky-University (Oldenburg, Germany) except *Phyllodoce empetriformis* (Sm.) D.D, collected in Botanical Garden Bochum (Bochum, Germany) with permission. For DNA extraction the leaves were silica gel dried, while for flow cytometry, we used fresh leaves. Vouchers for all species are stored in the herbarium of Carl von Ossietzky-University (OLD).

### Genome size estimation, DNA extraction and PCR amplification

To estimate nuclear genome sizes of *Rhododendron* species, we used flow cytometry following the basic protocol of Galbraith et al. (1983). We used *Zea mays* L. ‘CE-777’ (2C = 5.430 pg), *Hedychium gardnerianum* (2C = 4.02 pg) or *Solanum pseudocapsium* (2C = 2.60) as an internal standard (Temsch et al., 2010, Meudt et al., 2015; Table S2). Briefly, we prepared intact nuclei suspensions by chopping 0.5 - 1 cm^2^ of fresh leaf tissue of *Rhododendron* and the internal standard together with 1100μl of nuclei extraction buffer (OTTO I buffer: 100mM citric acid; 0.5 % (v/v) Tween 20 (pH approx. 2.3) after Otto (1990)). The intact nuclei suspension was filtered through a 30μm CellTrics^®^ nylon mesh filter (Partec). The filtered solution was then incubated at 37 °C for 30 min and stained with 2 ml propidium iodide buffer for 1 hour at 4 °C. This staining step also involved a treatment with RNase. Lastly, we ran the suspension on the flow cytometer (CyFlow SL, Partec, Munster, Germany), measuring 5000 particles and at least 1000 nuclei of sample and standard. We repeated this process three times on different days. The genome sizes (2C-value in pg DNA) were determined by comparing the mean relative fluorescence of each sample with the standard. The relationship between ploidy levels and genome sizes (monoploid 1Cx-value in pg DNA) was determined with documented chromosome numbers and ploidy levels in the literature.

We extracted total genomic DNA from silica gel-dried leaves (Chase and Hills, 1991), with minor modifications (100 μl elution buffer, centrifuge at 6.000 x g for 5 min) using the innuPREP Plant DNA Kit (Analytik Jena, Jena, Germany). Following upon the completion of DNA extraction, we amplified two plastid gene regions, *trn*K (*mat*K and the 3’ end *trn*K) & *trn*L-F (*trn*L intron, *trn*L 3’ exon, *trn*L-*trn*F spacer); and two nuclear gene regions, the *rpb*2-*i* (segment 2, 3, and 5) and the ribosomal Internal Transcribed Spacer (ITS) through polymerase chain reaction (PCR, Table S3). To amplify *trn*K, we designed two new internal primers based on the existing *trn*K of *Rhododendron setosum* D.Don (OLD00775) using the program PRIMER3 (Rozen and Skaletsky, 1999), i.e. MK1538F (TAT GGG TGT TTA AAG AGC) and MK1785R (TCT ATC ATT TGA CTC CGT ACC A). For other regions, we used the available primers (White et al., 1990; Taberlet et al., 1991; Liu et al., 1999). All amplification reactions of target regions were carried out in 25 μl with 2.5 μl 10x PCR buffer, 1 μl MgCl2 (50 mM), 0.5 μl dNTPs (10 mM), 1 μl of each primer (10 pmol/μl), 1 μl DMSO (only in case of nuclear regions), 1 μl BSA, 0.2 μl *Taq* polymerase (5 units), and 10-20 ng genomic DNA. The amplification of ITS and *trn*L-F regions were performed on a TProfessional Standard Thermocycler (Biometra GmbH, Goettingen, Germany) and those of the *trn*K and *rpb*2-*i* on a Mastercycler® gradient (Eppendorf AG, Hamburg, Germany; details of PCR reactions profile in Table S3). Sequencing was conducted by GATC Biotech (Konstanz, Germany) on an ABI 3730xl (PE Applied Biosystem) automated sequencers. To check the quality of all sequences, we used GENEIOUS PRO V5.4.6 (Kearse et al., 2012). Lastly, we used MAFFT algorithm (Katoh et al., 2002) to align the sequences and visually inspected the alignment.

### Phylogenetic reconstruction and divergence time

We analyzed the data considering them as four separate data sets (*trn*K, *trn*L-F, ITS, and *rpb*2-*i*) using the Maximum Likelihood and Bayesian inference approaches. Maximum Likelihood (ML) analyses were conducted in RAXML v.7.9.5 (Stamatakis, 2006) with the substitution model (GTR+Γ with four rate categories for ITS, *rpb*2 and *trn*K; and HKY for *trn*L-F). The selection of best substitution models was implemented in jMODELTEST v.3.7 (Posada, 2008) using the Akaike Information Criterion (AIC). We used non-parametric bootstraps (1000 replicates) to determine support for each node (BS up to 70 was considered as weak, BS 70 – 90 as medium, and BS > 90 as strong). Similarly, the Bayesian trees were analyzed in MRBAYES v.3.2.2 (Ronquist and Huelsenbeck, 2003) using the same substitution model with two independent runs, each consisted of four Markov chains. All runs were allowed to proceed for ten million generations, sampled every 1000 generations. A consensus tree was generated with a 50 percent majority rule consensus after discarding the first 10 percent generations as burn-in.

Similarly, we estimated time-calibrated phylogenetic trees in BEAST v.2.3 (Bouckaert et al., 2014) for which, the input file was generated in BEAUTi v.2.3 (Bouckaert et al., 2014) using an uncorrelated lognormal relaxed clock (Drummond et al., 2006) with a birth-death prior and GTR substitution model. Since fossils of *Rhododendron* are scarce and difficult to use for each branch, we used the oldest *Rhododendron* fossil as a first calibration point. According to Schwery et al., 2015, the estimated fossil age of *Rhododendron* (without *R*. *camtschaticum*) is 58 mya with a standard deviation of 2 my, based on the oldest fossil of *Rhododendron* and 17 other fossils from Ericaceae. As second calibration point, we used 28.10 mya as leaf fossil age of *R*. subgen. *Hymenanthes* from the late Oligocene (Axelrod, 1998) with normally distributed prior and the corresponding standard deviation.. The actual analysis was run for 100 million MCMC each, sampling the results after every 10.000 chains. We used the program Tracer v.1.5 (Rambaut and Drummond, 2009) to check upon the convergence of the chains and estimated sample sizes (ESS > 200). To compute the maximum clade credibility tree, we used TreeAnnotator v.2.3 (Drummond et al., 2012) with node heights being the median of the age estimates deleting the first 10 % generations as burn-in. Finally, we investigated if there were any influence of the priors on the analyses and information content of the data by repeating the analyses with the same settings without data.

### Genome size evolution and diversification

To investigate whether the continuous traits related to genome size (2C genome size, 1Cx monoploid genome size, ploidy level) have any significant phylogenetic signal, we estimated the phylogenetic signal of these characters with the function phylosig in the R-package ‘phytools’ (Revell, 2012) using two different methods: Blomberg’s *K* statistic (Blomberg et al., 2003) and Pagel’s λ (Pagel, 1999). A bar plot of the trait ‘ploidy level’ was visualized using ‘phytools’ by mapping it on the side of the BEAST tree. To calculate and map the ancestral character states for continuous characters 2C- and 1Cx-values on the BEAST tree for the data sets, we used the function contMap in ‘phytools’ by estimating the maximum likelihood ancestral character states for continuous traits with fastAnc.

The net diversification rate and the number and location of monoploid genome size rate shifts in *Rhododendron* were determined using BAMM (Rabosky, 2014) as employed in the R-package ‘BAMMtools’ (Rabosky et al., 2014). Though BAMM has been criticized by Moore et al. (2016), simulation studies suggested robustness of diversification analyses by BAMM (Rabosky et al., 2017; see also in Mitchel et al., 2019). The most important critique of BAMM is an error in the rate of extinction in the absence of fossil records (Rabosky, 2010; Marshall, 2017; and Rabosky, 2018). However, we here used BAMM only for diversification/speciation analysis. Additionally, since *Rhododendron* is monophyletic (details in results section), we only accounted for incomplete sampling to improve the robustness of BAMM based results. In *Rhododendron*, we estimated the globalSamplingFraction as number of species used in the dataset/total number of *Rhododendron* species (for example 0.235 for *trn*K as 259/1100 total *Rhododendron* species) following Igea and Tanentzap (2020; also see in Spriggs et al., 2015). For the actual analysis, we used the BEAST trees as input for running both the trait (1Cx-value) and speciation rate. Three replicates were run in BAMM for 30 million generations and saved after every 5,000^th^ generation. ‘BAMMtools’ was used to plot the likelihoods of sampled generations after discarding the first 10% of chains. To assess convergence, effective sample size was checked for each prior to be >200. Furthermore, we calculated Bayes factors (BFs), plotted the best shift rate configurations and estimated rates of speciation (diversification analysis) and evolution of monoploid genome size (trait analysis).

## RESULTS

### Molecular phylogenetic reconstruction

Details of variables sites, number of informative sites, and species used in each dataset are given in Table 1. The results from phylogenetic analyses of the four DNA regions (Figs. 1-2, S1-6) agree in several aspects. These are the monophyly of *Rhododendron*, the sister-group relationship of *R*. subgen. *Therorhodion* to the rest of the genus, the monophyly of *R*. subgen. *Rhododendron* (the former *Ledum* excluded) and *R*. subgen. *Tsutsusi* (except in *trn*LF), as well as the polyphyly of *R*. subgen. *Azaleastrum* and *R*. subgen. *Pentanthera*, within the latter, *R*. sect. *Pentanthera* is monophyletic if *R*. *canadense* is included (except in *rpb*2). In *R*. subgen. *Azaleastrum*, the two sections *R*. sect. *Azaleastrum* and *R*. sect. *Choniastrum* are monophyletic. Finally, the analyses agree that in *R*. subgen. *Tsutsusi, R*. sect. *Tsutsusi* (excl. *R*. *tashiroi*) and *R*. sect. *Brachycalyx* (including *R*. *tashiroi*) are monophyletic. In the following, we will only mention the points, in which the single region analyses differ from the rest.

**TABLE 1.** Numerical attributes of the four likelihood phylogenetic trees.

| Data set | *trnK* | *trnL-F* | ITS | *rpb2-i* |
| --- | --- | --- | --- | --- |
| No. individuals (Species) | 307(259) | 199(169) | 237(197) | 170(148) |
| Newly generated sequences | 204 | 119 | 162 | 82 |
| Aligned sequence length (bps) | 1750 | 983 | 660 | 3306 |
| No. of variable sites | 285 (16.30%) | 130 (13.20%) | 173 (26.20%) | 574 (17.30%) |
| No. of informative sites | 399 (22.80%) | 277 (28.10%) | 252 (38.10%) | 572 17.30%) |
| Consistency Index (CI) | 0.736 | 0.697 | 0.662 | 0.674 |
| Retention Index (RI) | 0.917 | 0.883 | 0.908 | 0.864 |

**Figure 1.**
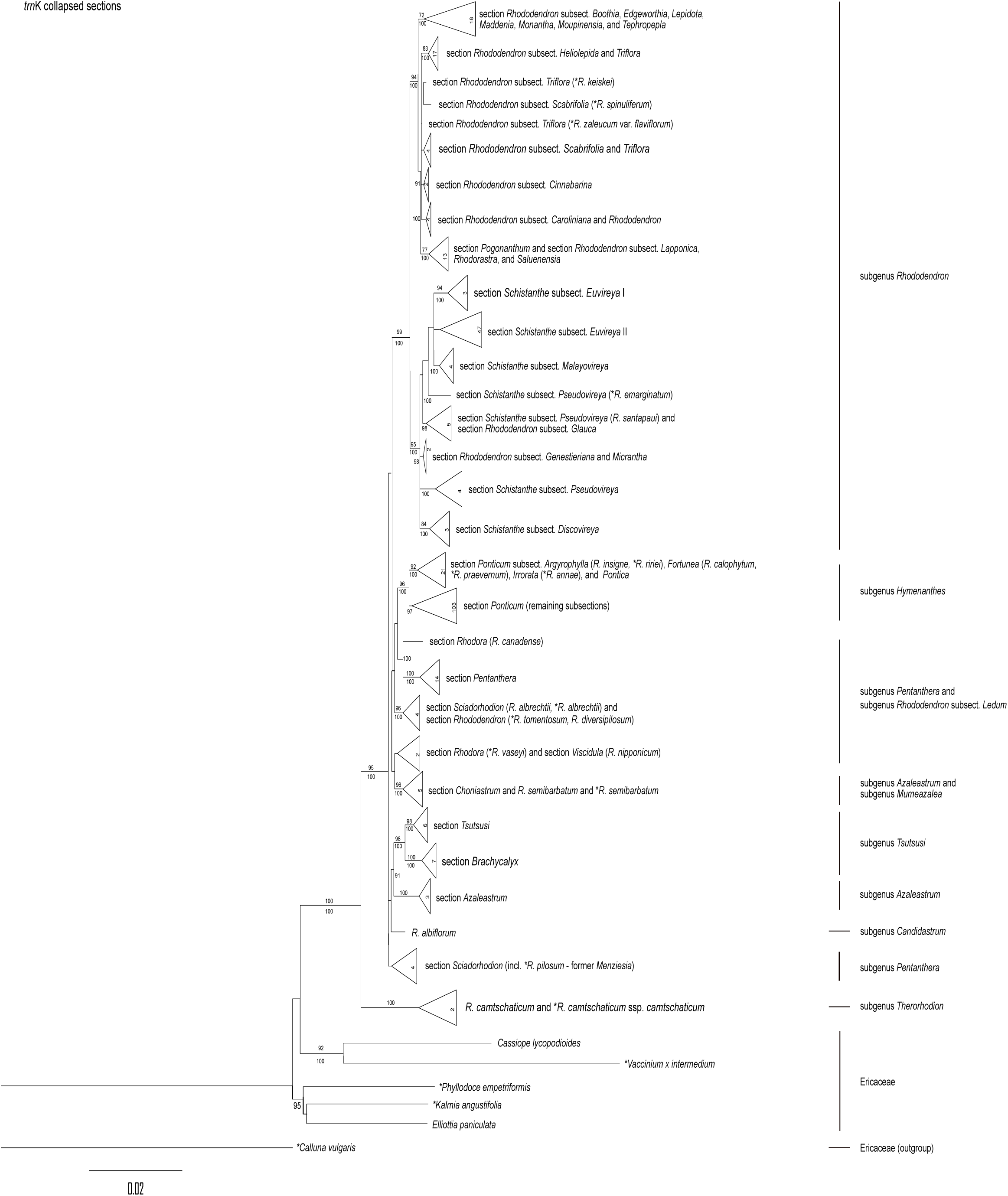
Phylogenetic tree of *Rhododendron* inferred from the plastid *trn*K gene region, showing species arranged by subgenus and section. Numbers above branches are bootstrap values, numbers below branches are posterior probabilities (in percent). Numbers in triangles are the number of the included individuals (species list in Table S2). Newly gener ated sequences are marked with a star (*).

Our results based on the plastid markers, the *trn*K phylogeny (259 species, CI = 0.736, RI = 0.917; Table 1; Fig. 1 and S1) and *trn*L-F region (169 species, CI = 0.697, RI = 0.883; Table 1; Fig. S2-3), are highly congruent. Results from the *trn*K dataset suggest that within *R*. subgen. *Hymenanthes*, species of subsections *Argyrophylla* (*R*. *insigne, R*. *rirei*), *Fortunea* (*R*. *calophytum, R*. *praevernum*), *Irrorata* (*R*. *annae*), and all included species of *Pontica* (except *R*. *degronianum* and *R*. *smirnowii*) form a group, which includes almost all species from outside SE Asia (South-West Eurasia = Turkey and Caucasus; North-East Asia = Japan, Korea and Manchuria to East Siberia; western North America and eastern North America) and is sister to the remaining subsections of *R*. subgen. *Hymenanthes* (SE Asian clade = mainly southern China, Himalaya Mountains and Taiwan). *Rhododendron* subgen. *Rhododendron* is divided into two clades of which one encompasses the sections *Rhododendron* (excluding subsection *Glauca*) and *Pogonanthum*, and the second contains the vireyas (section *Schistanthe*). The monotypic *R*. subgen. *Candidastrum* is sister to the combined *R*. subgen. *Tsutsusi* and *R*. section *Azaleastrum*, but the clade was weakly supported (BS/PP ≤ 70). The monotypic *R*. subgen. *Mumeazalea* is sister to *R*. sect. *Choniastrum*.

Our results based on the *trn*L-F region showed only some, slight differences and was generally less well resolved. The *trn*L-F-based tree recovered *R*. subgen. *Azaleastrum* section *Choniastrum* (with *Mumeazalea*) as sister to *Tsutsusi* section *Tsutsusi*. In addition, *R*. section *Azaleastrum* is sister to the clade of *R*. subgen. *Choniastrum* and *R*. subgen. *Tsutsusi* with strong support (Fig. S2). *Rhododendron vaseyi* (*R*. section *Rhodora*) and *R*. *pilosum* (*R*. section *Sciadorhodion*, former *Menziesia*) clustered closer to *R*. subgen. *Rhododendron* (Fig. S2). *Rhododendron* subgen. *Hymenanthes* is divided into the same two clades as above and within *R*. subgen. *Rhododendron* section *Schistanthe* (including *R*. section *Rhododendron* subsections *Genestieriana, Glauca*, and *Micrantha*) is sister to section *Pogonanthum* and the remaining subsections of section *Rhododendron*.

In the phylogeny based on the nuclear ITS region (197 species, CI = 0.662, RI = 0.908; Table 1; Fig. 2 and S4) reveals *R*. subgen. *Mumeazalea* clusters with section *Azaleastrum* and with *R*. section *Choniastrum*. In addition, *R*. sections *Rhodora* and *Sciadorhodion* show a sister group relationship to the clade of *R*. section *Azaleastrum, Mumeazalea*, and *Tsutsusi*, but this relationship is weakly supported. Within *R*. section *Rhododendron* the subsections *Ledum, Micrantha*, and *Rhododendron* form a clade (Fig. 2) which is sister to the clade including *R*. section *Pogonanthum*, the remaining subsections of *R*. section *Rhododendron*, and *R*. section *Schistanthe*.

**Figure 2.**
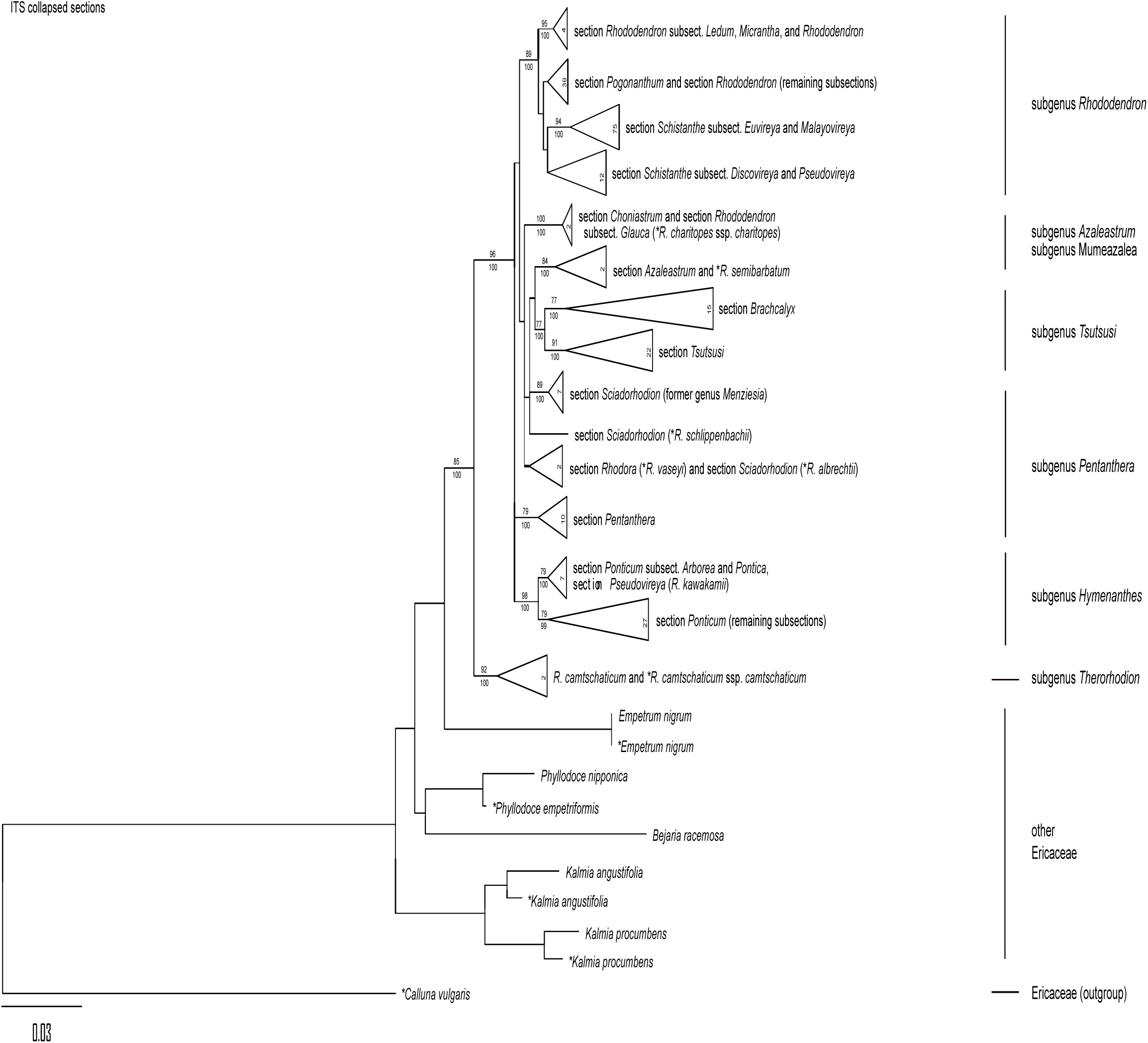
Phylogenetic tree of *Rhododendron* inferred from the nuclear Internal Transcribed Spacer (ITS) gene region, showing species arranged by subgenus and section*C*. *alluna vulgaris* (Ericaceae) was used as root. Numbers above branches are bootstrap values, numbers below branches are posterior probabilities (in percent). Numbers in triangles are the number of the represented individuals (species list in Table S2). Newly generated sequences are marked with a star (*).

In the phylogeny based on *rpb*2-*i* sequences (170 species, CI = 0.674, RI = 0.864; Table 1; Fig. S5-6) all *Rhododendron* species (except *R*. *camtschaticum*) fall into three large clades (Fig. S6). The first clade comprises *R*. subgenera *Hymenanthes* and *Pentanthera*, in which *R*. subgen. *Pentanthera* is not monophyletic. The second clade contains *R*. subgenera *Azaleastrum* section *Azaleastrum, Candidastrum, Mumeazalea, Pentanthera* section *Sciadorhodion, R*. *nipponicum* (section *Viscidula*) and *R*. *vaseyi* (section *Rhodora*), and *R*. subgen. *Tsutsusi*. The remaining clade encompasses *R*. subgen. *Azaleastrum* section *Choniastrum* and *R*. subgen. *Rhododendron*.

### Diversification regime shifts

All three runs of diversification analysis in BAMM showed similar results (including log likelihoods and number of shifts; data not shown). For the *trn*K species tree the frequent shift configuration of the 95 % credible set of shift configurations (f = 0.15) shows two ‘core shifts’ to higher diversification rates (red and orange circles and branches) and one ‘core shift’ to slower diversification rate (light blue circle and branches; Fig. 3). The first shift to diversification rate acceleration (red clade) includes species of *R*. subgen. *Hymenanthes* from the SE Asian clade. The second clade with higher diversification rate (orange clade) is within *R*. subgen. *Rhododendron*. This clade contains species from section *Rhododendron* in part (excluding species from *R*. subsect. *Baileya, Boothia, Edgeworthia, Lapponica, Maddenia*, and *Tephropepla*) and the two species included from *R*. section *Pogonanthum*. This diversification shift with rate acceleration is also found in the diversification analysis with only 105 taxa matching those included in the trait analysis (f = 0.35; Supplementary Material, Fig. S7A). The second shift within *R*. subgen. *Rhododendron* shows a diversification rate slowdown (light blue clade). The mean speciation rate in *Rhododendron* as calculated by BAMM was 0.429 speciation events per million years (Myr) and has increased over time (Fig. 3, inset).

**Figure 3.**
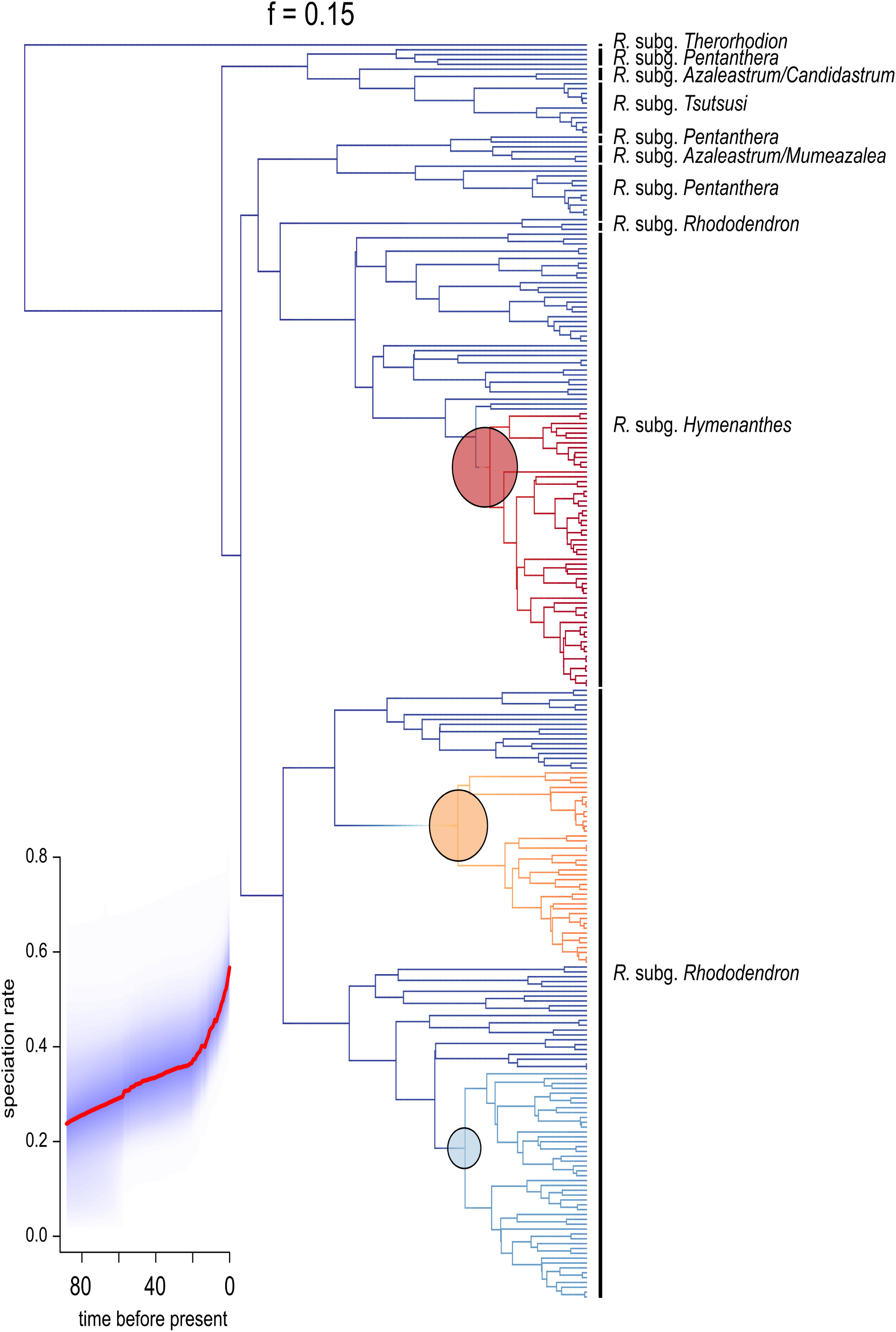
Result from the Bayesian Analysis of Macroevolutionary Mixtures (BAMM) diversification analysis of *Rhododendron trn*K. Colors of phylogeny refer to decrease (blue) or increase (red) in diversification rates. The size of the circles shows rate of diversification shift i.e., larger circle higher rate shift and smaller circle lowers rate shift. Inset shows the rate vs. time plot from the diversification analysis, where the ‘speciation rate’ (speciation events) is given per million years, and ‘time before present’ is in millions of years.

However, the diversification analysis of the *rpb*2-*i* species tree (f = 0.14) shows one ‘core shift’ to a higher diversification rate in *R*. subgen. *Hymenanthes* for the SE Asian clade (Fig. S8A) and a second ‘core shift’ with diversification rate slowdown for *R*. subgen. *Rhododendron* (excluding subsection *Ledum* (L.) K.A.Kron & W.S.Judd). Both shifts are not seen in the diversification analysis with only 56 taxa matching those for the trait analysis, despite a generally similar pattern (Fig. S8B). The diversification analysis of the ITS species tree shows only one ‘core shift’ from *R*. *camtschaticum* ssp. *camtschaticum* Pall. to a higher diversification rate for all remaining species of *Rhododendron* (Fig. S9A). In the diversification analysis with only 90 taxa a ‘core shift’ to a slower diversification rate for deciduous species (*R*. subgen. *Azaleastrum* section *Azaleastrum, R*. subgen. *Mumeazalea, R*. subgen. *Pentanthera* section *Sciadorhodion*, and *R*. subgen. *Tsutsusi* (Fig. S9B) has been revealed.

### Genome size evolution

The genome sizes for 125 *Rhododendron* species are listed in Table S2. The 1C-values range from 0.677 pg to 2.182 pg for *R*. subgen. *Hymenanthes*, from 0.543 pg to 1.914 pg for *R*. subgen. *Pentanthera*, from 0.483 pg to 2.777 pg for *R*. subgen. *Rhododendron* and from 0.571 pg to 0.776 pg for *R*. subgen. *Tsutsusi*. For *R*. subgen. *Azaleastrum* the genomes size is 0.583 pg and 1.406 pg, for *R*. subgen. *Mumeazalea* 0.540 pg and for *R*. subgen. *Therorhodion* it is 0.583 pg. Most (87%) species of *R*. subgenera *Hymenanthes, Pentanthera*, and *Tsutsusi* are diploid. In contrast, polyploid species (tetra-, hexa- and octoploids) constitute roughly half (53%) of all investigated species of *R*. subgen. *Rhododendron* sections *Rhododendron* and *Schistanthe*.

Among the three continuous traits (2C-value, 1Cx-value, and ploidy level), only the 1Cx-value had a significant phylogenetic signal (Blomberg’s K = 0.248, P < 0.005; Pagel’s λ = 0.929, P < 0.005). Similarly, the ancestral character state reconstruction analysis using continuous color gradients with 2C-values (Fig. 4A) indicates that the ancestors of *Rhododendron* had small genomes but there have been several increases (yellow to green/blue) of 2C-value along the tree (Fig. 4A). The species with larger genome sizes (green to blue) are mainly species of *R*. subgen. *Rhododendron* sections *Rhododendron* (in part) plus section *Pogonanthum* and section *Schistanthe*. In these two groups almost all polyploid species (87%) are included. The ancestral character state estimation of 1Cx-values indicates genome upsizing for species of *R*. subgen. *Pentanthera* section *Pentanthera, R*. subgen. *Hymenanthes*, and *R*. subgen. *Rhododendron* section *Rhododendron* subsect. *Baileya, Boothia, Edgeworthia, Maddenia*, and *Tephropepla* (green to blue). In contrast, species of *R*. subgen. *Rhododendron* section *Schistanthe* show a smaller monoploid genome size (red to yellow) which indicates genome downsizing for this group.

**Figure 4.**
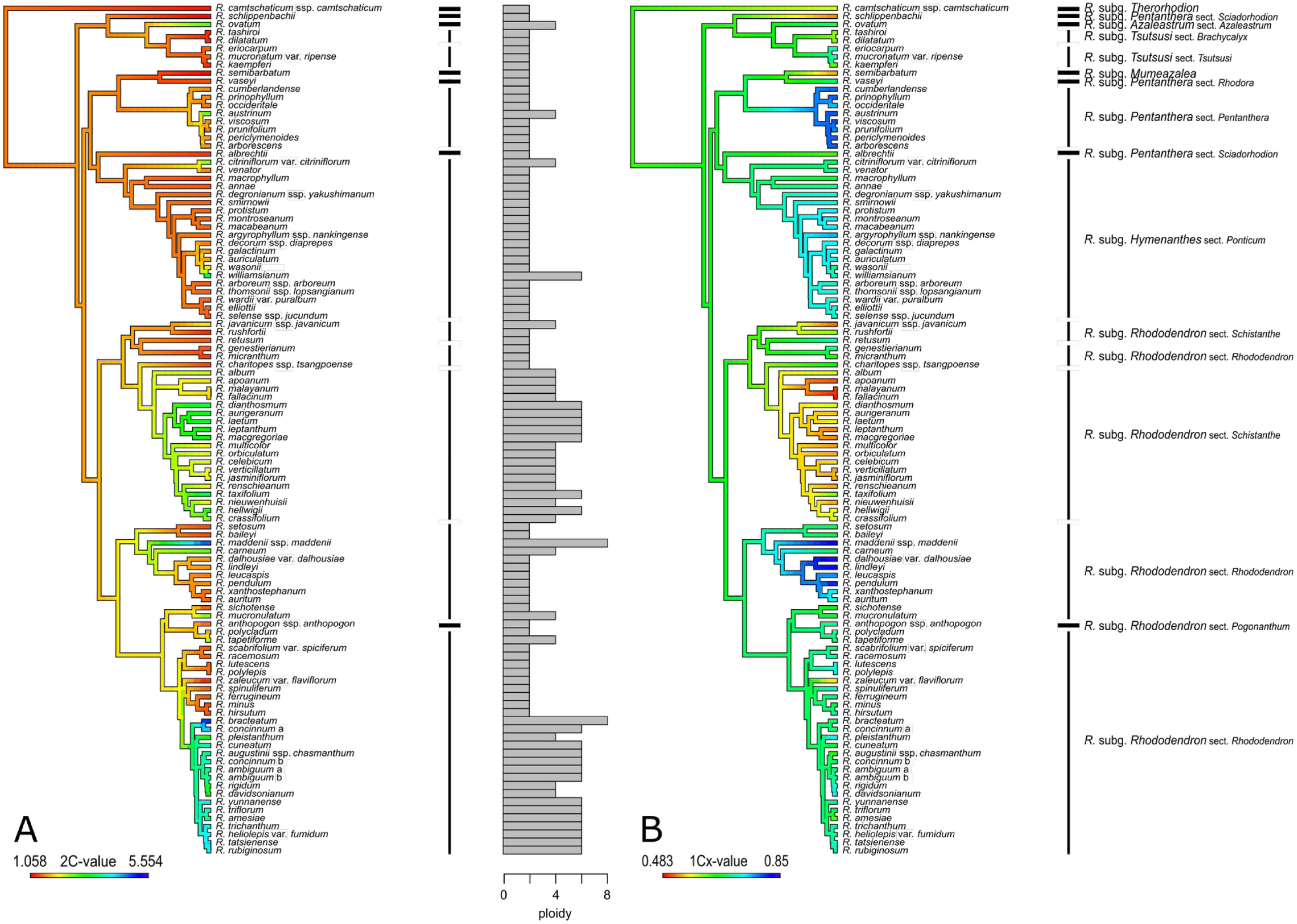
Ancestral genome size values mapped using a continuous color gradient on a BEAST tree of 105 Rhododendron species based on the trnK gene region. A, 2C-values and ploidy levels indicated as bar plot; B, 1Cx-values.

In the BAMM trait analysis of monoploid genome size (1Cx-value) the mean rate of monoploid genome size evolution in *Rhododendron* is 0.0005 and has not varied much over time (Fig. S10B, inset). Based on the *trn*K BEAST phylogeny three ‘core shifts’ are indicated in the trait analysis (Fig. S7B). Two of them are shifts to increased rates, i.e. in *R*. subgen. *Tsutsusi* section *Tsutsusi* and *R*. subgen. *Rhododendron* section *Malayovireya*, whereas a third shift to a decreased rate is shown for *R*. subgen. *Rhododendron* sections *Rhododendron* and *Pogonanthum*. Species of section *Rhododendron* subsect. *Baileya, Boothia, Edgeworthia, Maddenia*, and *Tephropepla* are excluded from this decrease.

The trait analysis based on the phylogeny of *rpb*2-*i* shows only one shift to an increased rate for *R*. subgen. *Tsutsusi* section *Tsutsusi* (Supplementary Material, Fig. S8C) and based on ITS only one ‘core shift’ to an increased rate for species of *R*. subgen. *Rhododendron* sections *Rhododendron* and *Pogonanthum* (except species of subsections *Micrantha* and *Rhododendron*; Fig. S9C) are indicated.

## DISCUSSION

Our results agree in a well-supported monophyly of *Rhododendron* (Figs. 1-2), which is in line with previous studies e.g., Shrestha et al., 2018; and Schwery et al., 2015), and when species of *Ledum* and *Menziesia* are included (Craven (2011), Goetsch et al. (2005), Kron and Judd (1990) and Kurashige et al. (2001). *Rhododendron camtschaticum*, representing *R*. subgen. *Therorhodion* is sister to the rest of the genus in all analyses. This agrees with other analyses and allows recognition of *R*. *camtschaticum* and *R*. *redowskianum* as separate genus (e.g., Judd and Kron, 2009). However, we continue recognizing them within *Rhododendron*.

In contrast, we found a marked difference concerning the internal relationships beyond the sectional level. Overall, the results revealed the larger a subgenus, the stronger is the support for its monophyly. The smaller subgenera pose several problems, for example *R*. subgen. *Azaleastrum* is inferred to be polyphyletic, as previously indicated (Goetsch et al., 2005; Kurashige et al., 2001; Schwery et al., 2015; and Yan et al., 2015). However, both sections are monophyletic in all analyses (inclusion of *R*. *charitopes* in the ITS analysis is dubious). However, several incongruent placements of species among analyses complicate a new classification of the genus. Shrestha et al. (2018) provided the most comprehensive phylogenetic analysis to date with multiple plastid and nuclear markers, but did not reflect incongruent placements among datasets (Table 2).

**TABLE 2.** Comparison of different molecular phylogenetic analyses with the results of this particular study. The abbreviated letters showed in the table represents, R – R. subgen. Rhododendron; A – *R*. sect. Azaleastrum; T – R. subgen. Tsutsusi; M – R. subgen. Mumazalea; C – R. sect. Choniastrum; S – R. sect. Sciadorhodion and P – R. sect. Pentanthera; H – R. subgen. Hymenanthes; L – R. sect. Ledum; V – R. sect. Viscidula.

|  | Kurashige et al. (2001) – *matK* | This study – *trnK* | This study – *trnLF* | Grimbs et al. (2017) – *trnK*, *trnLF*, ITS | Gao et al. (2002) – ITS | This study – ITS | Goetsch et al. (2005) – *rpB2* | This study – *rbB2* | Shrestha et al. (2018) – multiple regions |
| --- | --- | --- | --- | --- | --- | --- | --- | --- | --- |
| Sect. *Schistanthe* | Triphyletic (subsect. *Discovireya*/R. *santapaui*/rest) nested in R | unresolved within R | unresolved in R | - | single species nested in R | monophyletic nested in R | monophyletic *Discovireya* not sampled | diphyletic (Subsect. *Discovireya*/rest) nested in R | Diphyletic (subsect. *Discovireya*/rest) nested in R |
| *Ledum* | R. *hypoleucum* sister to R. *albrechtii* | R. *tomentosum*, R. *diversipilosum* sister to R. *albrechtii* | R. *tomentosum* close to R. *albrechtii* but sister to P | R. *tomentosum* sister to R. *albrechtii* | R. *tomentosum* sister to R | R. *tomentosum* sister to R | R. *hypoleucum*, R. *tomentosum* sister to R | R. *hypoleucum*, R. *tomentosum* sister to R | R. *hypoleucum*, R. *tomentosum* sister to R except R. *diversipilosum* sister to R. *albrechtii* |
| Sect. *Pentanthera* | sister to R. *canadense*, together sister to H | Sister to R. *canadense*, together sister to V, M, C, R. *vaseyi* | sister to R. *canadense*, together sister to L | sister to H (R. *canadense* not sampled) | sister to R. *canadense* in unresolved position | basally branching (R. *canadense* not sampled) | incl. R. *canadense* sister to H | together with R. *canadense* in H | incl. R. *canadense*; sister to H |
| Sect. *Choniastrum* | sister to M | sister to M | sister to M | sister to T, A | unresolved with A, T, R. *albiflorum*, R. *schlippenbachii* | sister to A, T, S, M, R. *vaseyi* | sister to R, L | sister to R, L | sister to R, L |
| R. subgen. *Tsutsusi* | sister to A | sister to A | paraphyletic sister to A+M | sister to A | sister to A+M | sister to A+M | unresolved with A, M, V | sister to V, together sister to M, together sister to A | sister to A+M |
| R. sect. *Azaleastrum* | sister to T | sister to T | sister to T, C, M | sister to T | Sister to M | sister to M | unresolved with T, M, V | unresolved with T, M, V | sister to T, M |
| R. *albiflorum* (R. subgen. *Candidastrum*) | unresolved with T, A, R. sect. *Sciadorhodion* | sister to T, A | - | - | Unresolved with T, A, C, R. sect. *Sciadorhodion* | - | sister to R. *albrechtii* | - | sister to R. *albrechtii* + R. *diversipilosum* |
| R. *semibarbatum* (R. subgen. *Mumeazalea*) | sister to C | sister to C | sister to C | - | sister to A | sister to A | sister to V | sister to V, T, together sister to A | sister to T |
| R. *nipponicum* (R. sect. *Viscidula*) | sister to M, C | sister to R. *vaseyi* | - | - | - | - | sister to M | sister to T | Sister to R. *quinquefolium* (sect. *Sciadorhodion*), together sister to A, T, M |
| R. *vaseyi* (R. sect. *Rhodora*) | - | sister to V | sister to R | sister to R, H | - | sister to R. *albrechtii* | sister to *Menziesia*, together sister to R. *schlippenbachii* | sister to R. *schlippenbachii*, together sister to *Menziesia* | with R. *pentaphyllum* sister to *Menziesia*, together sister to R. *schlippenbachii* |
| *Menziesia* | Unresolved with A, T, C, R. sect. *Sciadorhodion* | R. *pilosum* incl. in R. sect. *Sciadorhodion* | R. *pilosum* with R. *vaseyi* sister to R | - | - | 5 spp. sister to R. *schlippenbachii* | 2 spp. sister to R. *vaseyi*, together sister to R. *schlippenbachii* | - | 9 spp. sister to R. *vaseyi* and R. *pentaphyllum*, together sister to R. *schlippenbachii* |
| R. *schlippenbachii* (R. sect. *Sciadorhodion*) | unresolved with *Menziesia*, A, T, R. *pentaphyllum*, R. *quinquefolium* | together with *Menziesia* and R. *pentaphyllum*, R. *quinquefolium*, close to A, T, V. *albiflorum* | unresolved | basally branching | unresolved with T, A, R. *albiflorum* | sister to *Menziesia*, together sister to T, A, M | sister to R. *vaseyi* + *Menziesia*, then to R. *albiflorum*, then to A, T, M, V | sister to R. *vaseyi*, together sister to *Menziesia*, sister to A, T, M, V | sister to *Menziesia* + R. *vaseyi* |
| R. *albrechtii* (R. sect. *Sciadorhodion*) | sister to R. *hypoleucum* (L), together unresolved | R. *albrechtii* sister to L, together sister to H | unresolved | sister to L, together sister to P, H | - | sister to R. *vaseyi*, together sister to C | R. *albrechtii* sister to R. *albiflorum* | sister to R. *vaseyi*, R. *schlippenbachii* + *Menziesia* | sister to R. *diversipilosum* (L), together sister to R. *albiflorum* |

### Phylogeny of *Rhododendron*

Goetsch et al. (2005) grouped the species in five subgenera, *R*. subgen. *Therorhodion* with two species and *R*. subgen. *Rhododendron, Hymenanthes, Azaleastrum* and *Choniastrum* (Table 2). The largest of these subgenera is *R*. subgen. *Rhododendron* with more than 500 species (Table S1). There have been debates on whether to recognize the vireyas as separate subgenus or section (Craven et al., 2008; Argent and Twyford, 2012). Recognition of a separate subgenus in the traditional sense would lead to recognition of a diphyletic clade since *R*. subsection *Discovireya* is separate from the rest in most analyses as found earlier by Goetsch et al. (2011). The phylogeny presented by Shrestha et al. (2018) would allow recognition of three subclades, one containing most of the members of *R*. sect. *Schistanthe*. However, these clades are inconsistent among studies and markers and we refrain from suggesting a new sectional classification. The only question regarding the circumscription of this subgenus is the inclusion of the former genus *Ledum*, which appears as sister to *R*. subgen. *Rhododendron* in all nuclear DNA-based analyses (Fig. 2 and S1; Gao et al., 2002; and Goetsch et al., 2005) but as sister to *R*. *albrechtii* (Fig. 1 and S5; Kurashige et al., 2001) or sister to *R*. subgen. *Hymenanthes* and *R*. sect. *Pentanthera* (Schwery et al., 2015) and distant to *R*. subgen. *Rhododendron* in the plastid DNA-based analyses. Shrestha et al. (2018) has some species as sister to *R*. subgen. *Rhododendron* and some as sister to *R*. *albrechtii* in his combined analysis of plastid and nuclear DNA, which suggests that missing data in their dataset causes different accessions of species from this group to end up in different positions. In Grimbs et al. (2017) the plastid DNA signal apparently overruled the nuclear signal.

The second largest group in *Rhododendron* is *R*. subgen. *Hymenanthes*. Whereas there is mostly strong support for the morphologically well-circumscribed subgenus in the traditional sense (except in *rpb*2; Fig. S5-6; Goetsch et al., 2005), Shrestha et al. (2018) enlarged the subgenus to include *R*. sect. *Pentanthera*. This relationship is strongly supported by *rpb*2 (Fig. S5-6; Goetsch et al., 2005) and shown without support by *trn*K (Fig. 1; Kurashige et al., 2001; Schwery et al., 2015). The relationship is not shown but also not strongly refuted by analyses of ITS (Fig. 2 and S4; Gao et al., 2002). The two groups differ in leaves being either deciduous or evergreen, a character traditionally in taxonomy and horticulture important in the distinction between rhododendrons and azaleas. We, therefore, prefer to keep *R*. sect. *Pentanthera* separate from *R*. subg. *Hymenanthes*.

The second-smallest subgenus in the classification of Goetsch et al. (2005) is *R*. subg. *Choniastrum* with 15 species. The signal in Shrestha et al. (2018) depicting it as sister to *R*. subgen. *Rhododendron* is largely derived from the *rpb*2-dataset (Fig. S5-6; Goetsch et al. 2005). However, ITS puts the group in a position as sister to *R*. subg. *Azaleastrum* sensu Goetsch et al. (2005; Fig. 2) or in unresolved position in one clade with this subgenus (Gao et al., 2002). Schwery et al. (2015) resolves the section including former genus *Diplarche* based on *mat*K and *rbc*L as sister to this clade, as well. Other studies based on plastid DNA markers even group the section among the members of *R*. subgen. *Azaleastrum* sensu Goetsch et al. (2005). We, therefore, consider it premature to recognize this clade at the subgeneric level.

The morphologically most heterogeneous subgenus of Goetsch et al. (2005) is *R*. subgen. *Azaleastrum*, which includes *R*. sect. *Azaleastrum*, three sections formerly assigned to *R*. subgen. *Pentanthera* (*R*. sect. *Rhodora, R*. sect. *Sciadorhodion* (incl. *Menziesia*), *R*. sect. *Viscidula*), the monotypic *R*. subgen. *Mumeazalea* (*R*. *semibarbatum*) and *R*. subgen. *Candidastrum* (*R*. *albiflorum*) and the large (66 species) *R*. subgen. *Tsutsusi*. The signal for the monophyly of this clade is derived mostly from *rpb*2 (Fig. S5-6; Goetsch et al., 2005) with ITS supporting it with inclusion of *R*. sect. *Choniastrum* (Fig. 2 and S4; Gao et al., 2002). Plastid DNA does not support the group as monophyletic. The core of the group is constituted by *R*. subgen. *Azaleastrum* and *R*. subgen. *Tsutsusi*. The monotypic *R*. subgen. *Candidastrum* (*R*. *albiflorum*) clusters with these two based on *trn*K (Fig. 1 and S1; Kurashige et al., 2001), is unresolved together with these based on ITS (Gao et al., 2002) and strongly supported sister to *R*. *albrechtii* based on *rpb*2 (Goetsch et al., 2005) and in Shrestha et al. (2018). The case is the other way around for the monotypic *R*. subgen. *Mumeazalea* (*R*. *semibarbatum*), which is sister to *R*. subgen. *Tsutsusi* in Shrestha et al. (2018) or *R*. subgen. *Azaleastrum* with ITS (Fig. 2 and S4; Gao et al., 2002) or in one clade with these two and *R*. sect. *Viscidula* in *rpb*2 (Fig. S2-3; Goetsch et al., 2005) but distantly to those and strongly supported sister to *R*. sect. *Choniastrum* in the plastid DNA-based phylogenies (Fig. 1 and S2-3; Kurashige et al., 2001). A third species (group) related to this core group is mono- or ditypic *R*. sect. *Viscidula* (*R*. *nipponicum*), which clusters with this core in *rpb*2 (Fig. S5-6; Goetsch et al., 2005) and in Shrestha et al. (2018) but not using *trn*K (Fig. 1; Kurashige et al., 2001) with weak support. A fourth species changing positions in different analyses is *R*. *vaseyi*, the second species of *R*. sect. *Rhodora* apart from *R*. *canadense*, which clusters with *R*. sect *Pentanthera*. *Rhododendron vaseyi* is related to *R*. *schlippenbachii* of *R*. sect. *Sciadorhodion* and the species of former genus *Menziesia* with medium to strong support based on *rpb*2 (Fig. S5-6; Goetsch et al., 2005) and is found there in Shrestha et al. (2018), too. However, ITS and the plastid DNA markers retrieve it in different positions (Fig. 1-2 and S1-6; Kurashige et al., 2001; Grimbs et al., 2017). The former genus *Menziesia* has been found to be nested in *Rhododendron* since the studies of Kron (1997) and Kurashige et al. (2001). Goetsch et al. (2005) found strong support for a clade consisting of *Menziesia* with *R*. *schlippenbachii* and *R*. *vaseyi* and a relationship with either holds in all analyses but not always with both but almost always with *R*. *schlippenbachii*. In turn, *R*. *schlippenbachii* (and *Menziesia*) have different positions, but always close to the Azaleastrum-Tsutsusi-group. It is, however, noteworthy that other members of *R*. sect. *Sciadorhodion* rarely form a monophyletic group with these. For example, *R*. *albrechtii*, which takes only a distant relationship with *R*. *schlippenbachii* in the plastid DNA-based analyses (Fig. 1 and S1-3; Kurashige et al., 2001; Grimbs et al., 2017).

Taken together, we consider it premature to group all species of *Rhododendron* in five subgenera. We see the merit in the recognition of the five major clades at the subgeneric level but given the amount of incongruence a large number of species cannot be confidently assigned to one of these five clades. Further, genome-wide data will be necessary to assess whether these currently unassignable taxa are independent taxa, assignable to one of the five major clades or whether they are inter-subgeneric hybrids.

### Diversification regime shifts

Based on this incongruence, we considered it necessary to conduct diversification analyses separate for each DNA marker, although the *trn*LF-region did not have enough variation to allow a reliable analysis. Diversification analyses demonstrated nearly the same speciation shifts for the plastid *trn*K and nuclear *rpb*2-*i* regions (Fig. 3 and Fig. S8). The main shift to higher diversification was found for species of *R*. subgen. *Hymenanthes* from the Himalayan/SE Asian clade, although the exact species included is not consistent. Most species of subsection *Pontica* with a distribution outside SE Asia (e.g., SW Eurasia, NE Asia, and N America) show no shift in diversification rate. This pattern is consistent with the findings of other analyses. For example, Milne et al. (2010) hypothesized, based on divergence time estimations, a slow diversification outside SE Asia followed by more rapid diversification of one lineage within SE Asia, followed by immigration of at least one additional lineage to the region. Additional support for a nested Himalayan radiation is given by Schwery et al. (2015) using a BAMM analysis similar to ours but with much smaller taxon sampling. The Himalayan-Southwest China region is known as a species-rich area of *Rhododendron* since Joseph D. Hooker famous travels to India and the Himalayas (1847 – 1851) but the region is also well known for other highly diverse groups of plants (Qiu et al., 2011). Most intriguing is the parallel increase in diversification in shrubby *Viburnum* of the region, which has been dated to the Eocene (Spriggs et al., 2015), similar to *Rhododendron*. Suggested reasons for the high diversity that may apply to *Rhododendron* are the climatic and physiographic heterogeneity, a complex geological history and the absence of major Quaternary glaciations (Qiu et al., 2011) coupled with an increase in precipitation (Wang et al., 2012). These factors may have spurned a diversification by causing barriers to plant migration (Zhao et al., 2013) and providing opportunities for frequent niche shifts between temperate and tropical biomes with limited vertical migration (Spriggs et al., 2015).

Similar to *R*. subgen. *Hymenanthes*, a significant rate increase was found in most diversification analyses in *R*. subgen. *Rhododendron* (Figs. 3 and Fig. S7) and comprises most species of sections *Pogonanthum* and *Rhododendron*. In contrast, a clade within section *Rhododendron* containing species of section *Rhododendron* subsections *Baileya, Boothia, Edgeworthia, Maddenia*, and *Tephropepla* shows no change in diversification rate. All members of this clade without rate shift occur in the Himalayan Mountains above 1500 meters with many at least facultative epiphytic species and in an ecological diverse array of habitats such as conifer forests, grassy hillsides, among rocks, steep slopes, or cliffs. In contrast to our expectations, no shift in diversification rate was found within the species-rich group of tropical *Rhododendron* (vireyas, section *Schistanthe*) for subsection *Euvireya* (Fig. 3; Figs. S7-8) which includes almost all species endemic to the Philippines, Borneo, New Guinea, Sulawesi, and the Solomon Islands. Nevertheless, here we only included 22% species of *Rhododendron* out of their total 587 species (Table S1). *Euvireya* species exhibit considerable variation in their ecology, being epiphytic or terrestrial and occurring from sea level to over 4000 m. Within *Euvireya*, the molecular-phylogenetic clustering follows geography more closely than traditional taxonomy based upon morphology, which is similar to the results from (Goetsch et al., 2011). The geographical pattern of the group was intensively discussed by Brown et al. (2006b). It is considered a classic example of Malayan radiation (Brown et al., 2006b; and Goetsch et al., 2011). An adaptive radiation can be accompanied by a diversification rate slowdown, which may sound counter-intuitive at first. However, since speciation rates usually decrease after an initial rapid diversification either due to increased competition for resources or niche filling, diversity- or time-dependent factors must be considered as potential drivers for observed decreases in the rates of diversification (Soulebeau et al., 2015). Our results are in contradiction with Shrestha et al. (2018), who suggested that tropical and subtropical mountains are not only the biodiversity and endemism hotspots for the genus *Rhododendron*, but also function as cradles of *Rhododendron* diversification.

While support for a diversification increase in *R*. subgen. *Hymenanthes* is more or less unambiguous, we have found here a discordance of results from different DNA regions but cannot really distinguish between diversification rate shifts depending on species sampling, evolutionary rate or topology (Fig. 3; and Figs. S7 and S9). Based on the similarity of our analysis using *trnK* with 260 taxa and that of Schwery et al. (2015) with 60 taxa but also using *trn*K (and *rbc*L), topology seems to be the most important. Therefore, more complete taxon sampling and, especially, resolving incongruences between markers in the future seems to be required for more conclusive diversification analyses.

### Genome size evolution

Our analysis is the first to analyze genome sizes in a larger number of species of *Rhododendron* (Table S2). Previous genome size estimations had been generated using Feulgen densitometry (Ammal, 1950; Ammal et al., 1950) or flow cytometry using DAPI staining (Jones et al., 2007). Ammal et al. (1950) completed an extensive survey of chromosome numbers and ploidy levels in *Rhododendron* and found the elepidote rhododendrons (*R*. subgen. *Hymenanthes*), evergreen azaleas (*R*. subgen. *Tsutsusi*), and the deciduous azaleas (*R*. subgen. *Pentanthera*) to be predominantly diploid. However, they also demonstrated the occurrence of triploids, hexaploids, octoploids, and dodecaploids (2n = 12x = 156) within *R*. subgen. *Rhododendron* and natural tetraploids in other subgenera as well, e.g. in *R*. subgen. *Pentanthera* (*R*. *canadense* and *R*. *calendulaceum;* Ammal, 1950; Ammal et al., 1950; and Jones et al., 2007). Our results confirm and expand the pattern that polyploidy occurs in almost all subgenera but most polyploid species are within *R*. subgen. *Rhododendron* sections *Rhododendron* and *Schistanthe* (Fig. 4A). This is rather surprising given the high frequency of polyploids among garden cultivars belonging to *R*. subgen. *Hymenanthes* (Perkins et al., 2012). Within *R*. section *Schistanthe*, the polyploid species are restricted to *R*. subsections *Euvireya* and *Malayovireya*, which occur in Indonesia and New Guinea and most of them are endemic to these islands while the species of the Asian mainland are diploid based on our data. There seems to be no correlation of genome size with habit (epiphytic, terrestrial or both) in this group of vireyas but the monoploid genome sizes are smaller (1Cx-values = 0.483 up to 0.618 pg; light green/yellow to red; Fig. 4B). This genome downsizing has been inferred to be accompanied by a slow-down in the evolution of genome size. Thus, the group seems to have stabilized on a lower level of genome size. Such a pattern has also been shown in the New Zealand radiation of *Veronica*, in which genome downsizing and a slow-down of rate in genome size evolution are associated with a radiation on the polyploid level (Meudt et al., 2015). While previous studies have shown that genome sizes of tropical angiosperms in general are lower than those of colder, temperate regions (Levin and Funderburg, 1979; Ohri, 2005), our study seems to be the first to indicate such a shift to small genome-tropical clade within a specific genus. It remains to be studied whether genome downsizing in this clade is functionally related with the higher frequency of polyploidy and/or its species richness.

Within section *Rhododendron*, almost all polyploid species form a monophyletic clade and seem to be restricted to subsections *Heliolepida* and *Triflora*. In contrast to the former group, here polyploidy is not associated with genome downsizing. Diploid species from the latter subsection do not cluster with the monophyletic clade of polyploids but with other diploid species from different subsections based on plastid DNA markers (Fig. 4A; *trn*K gene region). In the same analysis using ITS (Fig. S10) diploids are nested among polyploids suggesting an origin of at least four polyploid clades involving the same unknown diploid species. Hybridization and polyploidization, often referred to as whole genome duplication, are both potential speciation mechanism. Genome doubling creates instantaneously lineages reproductively isolated from its diploid progenitors and must overcome competition with their parents for abiotic resources (Soltis et al., 2010). Diversification analyses have demonstrated that polyploids have higher extinction rate and lower diversification rate than diploid lineages (Mayrose et al., 2011) but nevertheless have given rise to major radiations in angiosperms (Tank et al., 2015). In line with this pattern, we did not detect a change in diversification in polyploids. Thus, polyploidy did not increase speciation rate in *Rhododendron* but may however be associated with evolutionary novelties. Our hypothesis is that in this Heliolepida/Triflora-group species have a higher chance to produce unreduced gametes, thus the repeated origins of polyploids from similar ancestors, but the lack of genome downsizing prevents diversification on the polyploid level.

Hybridization facilitates the transfer of traits between species, which has been shown to promote adaptive evolutionary change in *Rhododendron* (Milne and Abbott, 2000) and other species. Such introgressed traits may affect various stages of life history including resistance to herbivores and pathogens (Whitney et al., 2015). Recent reviews suggest that resistance is an important component of hybrid survival (Orians, 2000) and that hybridization and polyploidy may be important evolutionary mechanisms for generating novel secondary chemicals important in the diversification of plant-animal interactions (Soltis et al., 2014). Oswald and Nuismer (2007) explored the possibility that new polyploids are initially more resistant to pathogens than their diploid progenitors by using mathematical models and confirmed that polyploids are significantly more resistant. *Rhododendron* may prove to be a living example for the evolution of novel chemicals in polyploids since four of the ten species shown to exhibit the highest antibacterial effects against Gram-positive bacteria by Rezk et al. (2015) are shown here to be polyploid (*R*. *ambiguum, R*. *cinnabarinum, R*. *concinnum* and *R*. *rubiginosum*). It remains to be shown that the high antimicrobial activity is due to the origin of novel gene combinations in polyploids.

## CONCLUSIONS

Given the large numbers of species, including many rare species with restricted range sizes, there are inherent difficulties in the analyses of phylogenetic relationships, diversification, and trait evolution. Here, we included only 25-30% of the total species from the genus in the study and analyzed all data sets separately. However, by using state of the art analytical tools, relying on strong support in terms of bootstraps and posterior probabilities, and discussing thoroughly and critically the results, we provide the most up-to-date knowledge of *Rhododendron* phylogeny, diversification and genome size evolution. We compared our results with the DNA-based classification by Goetsch et al. (2005) and warn of adopting this classification uncritically (Table 2). Our results agree in a well-supported monophyly of *Rhododendron*, with *R*. subgen. *Therorhodion* sister to all other taxa of *Rhododendron*. The results recovered marked incongruences between markers in retrieval of internal relationships but also finding many results common across DNA markers and morphology-based classifications. Our results demonstrate that the definition of enlarged *R*. subgen. *Azaleastrum* and *R*. section *Sciadorhodion* by Goetsch et al. (2005) including *R*. subgen. *Candidastrum, R*. subgen. *Mumeazalea, R*. *vaseyi* and other species of *R*. sect. *Sciadorhodion* may be premature. Similarly, our results do not consistently support the placement of *R*. subsect. *Ledum* in *R*. subgen. *Rhododendron* and the inclusion of *R*. sect. *Pentanthera* in *R*. subgen. *Hymenanthes*. The diversification analysis revealed that a major rate shift in *Rhododendron* occurred in the Himalayan Mountains above 1500 meters with many at least facultative epiphytic species and in an ecologically diverse array of habitats such as conifer forests, grassy hillsides, among rocks, steep slopes, or cliffs. These results are in contradiction to Shrestha et al. (2018), who suggested tropical and subtropical mountains to be not just the biodiversity and endemism hotspot for the genus but also a cradle of its diversification. However, the topology of the phylogeny is indicated to influence largely the results. Therefore, more complete taxon sampling, especially, resolving incongruences between markers in the future seems to be required for more conclusive diversification analyses. Lastly, we confirm and expand that polyploidy occurs in almost all subgenera but most polyploid species are within *R*. subgen. *Rhododendron* sections *Rhododendron* and *Schistanthe*. The two groups differ, however, in their pattern with the polyploids in *R*. sect. *Rhododendron* originating frequently but not diversifying on the polyploid level and not exhibiting genome downsizing. In contrast, *R*. sect. *Schistanthe* seems to be another example for a polyploid radiating after genome downsizing. It further can serve as an example for the frequently cited reduction in genome size of tropical plants versus their temperate relatives.

## ACKNOWLEDGEMENTS AND FUNDING INFORMATION

This study was financially supported by the Stiftung Bremer Rhododendronpark, Germany. The authors would like to thank Silvia Kempen and Eike Mayland-Quellhorst for assistance in the lab, Nikolai Kuhnert, Klaudia Brix and Matthias Ullrich for support of this project.

## DATA ACCESSIBILITY

Postprocess data will be available on Dryad.

## AUTHOR CONTRIBUTIONS

Conceived the idea: DCA, JN, HS and KG; Performed experiments: JN; Contributed reagents/materials/analysis tools: DCA, and HS; Analyzed data and authored the drafts of the paper: KG, DCA. All authors contributed, made multiple revisions, and approved the final draft.

