## Supplementary figures and images for "Incongruent phylogenies and its implications for the study of diversification, taxonomy and genome size evolution of *Rhododendron* (Ericaceae)"

### Supplemental Figure 1

*trn*K full tree

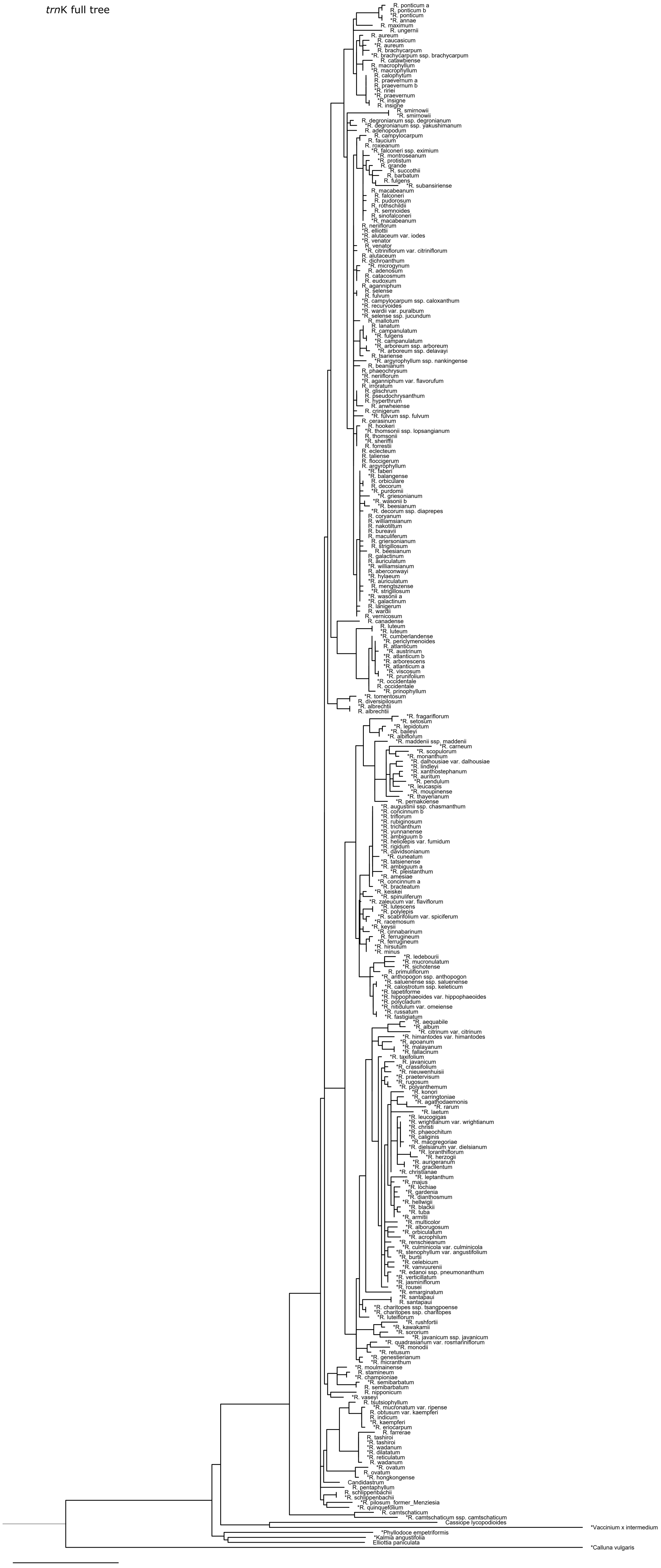

### Supplemental Figure 2

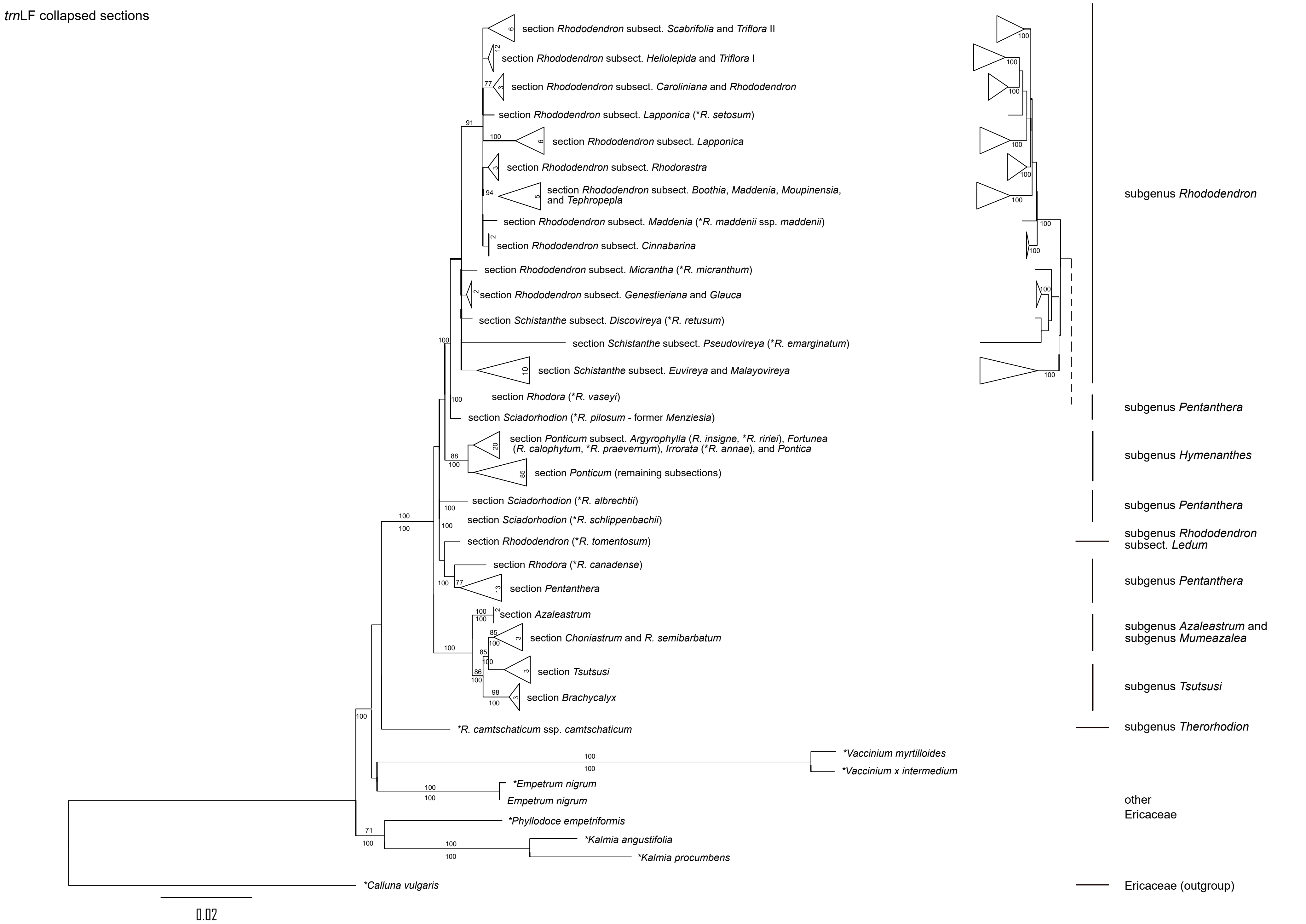

### Supplemental Figure 3

*trn*L-F full tree

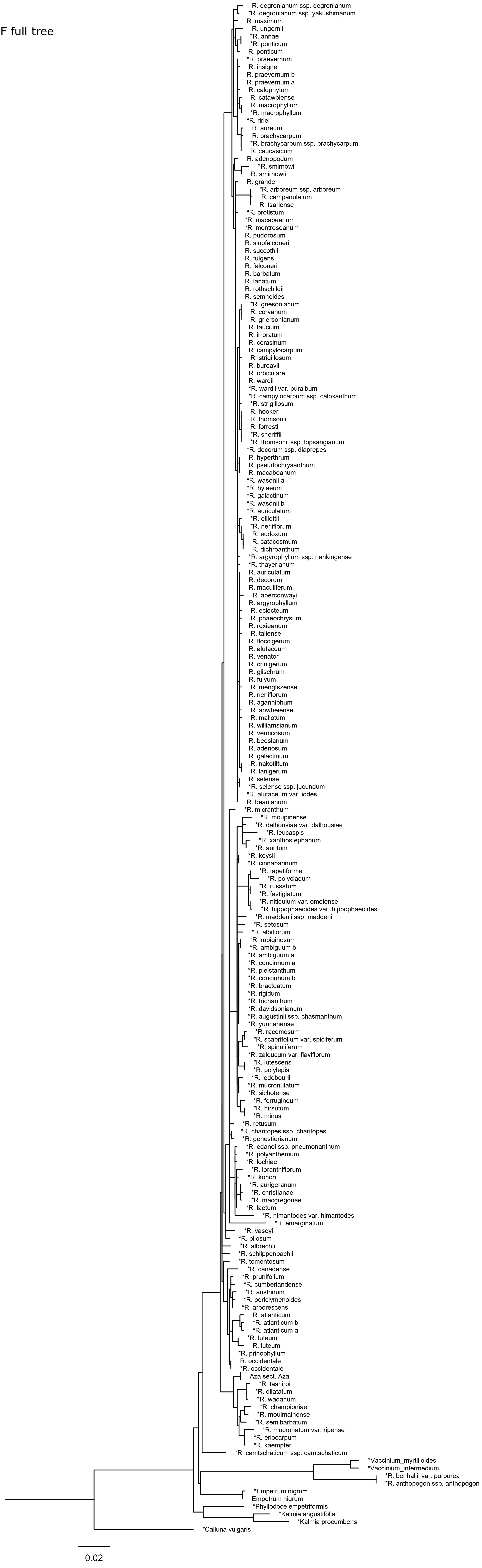

### Supplemental Figure 4

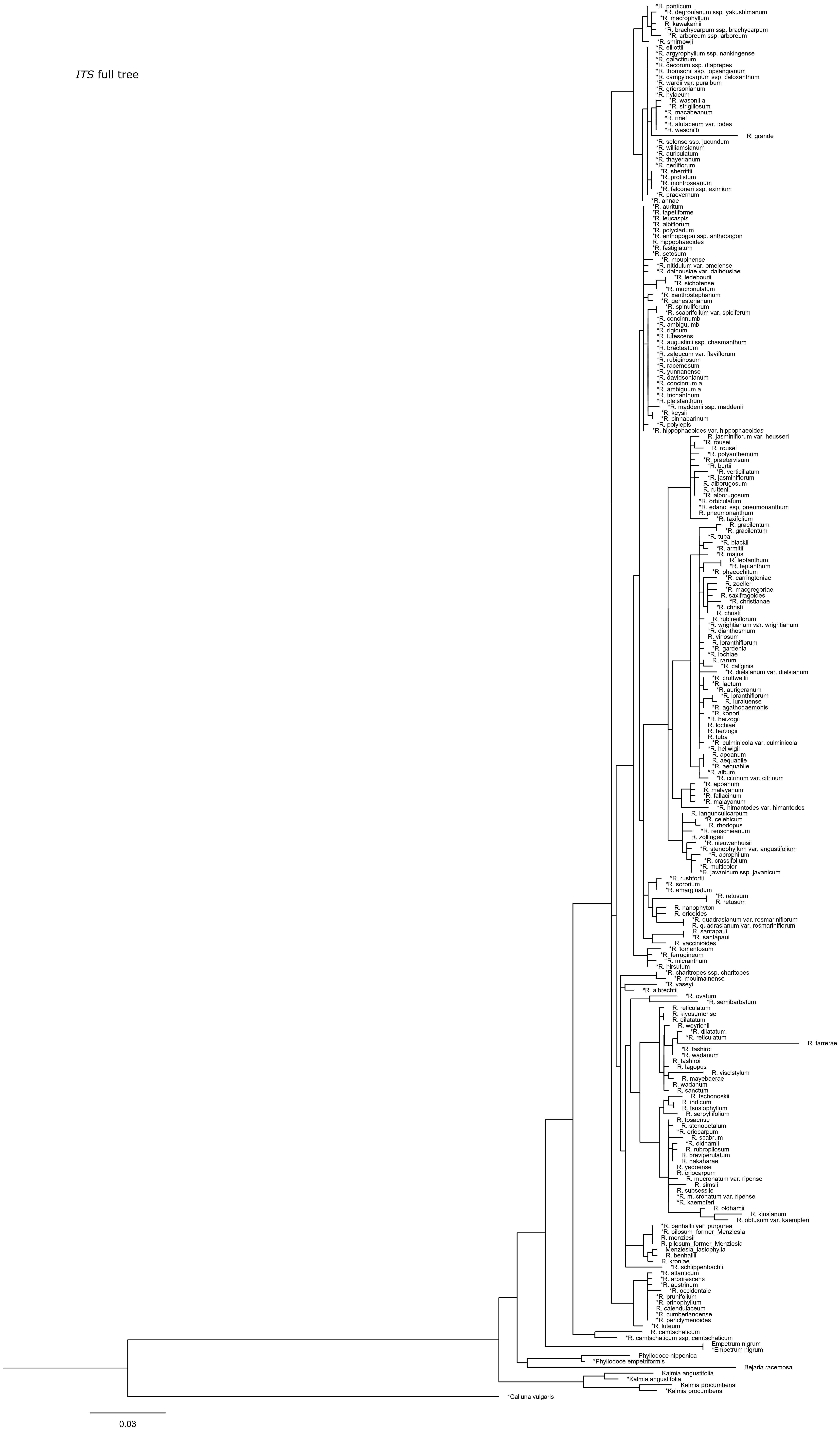

### Supplemental Figure 5

rpb2-i collapsed sections

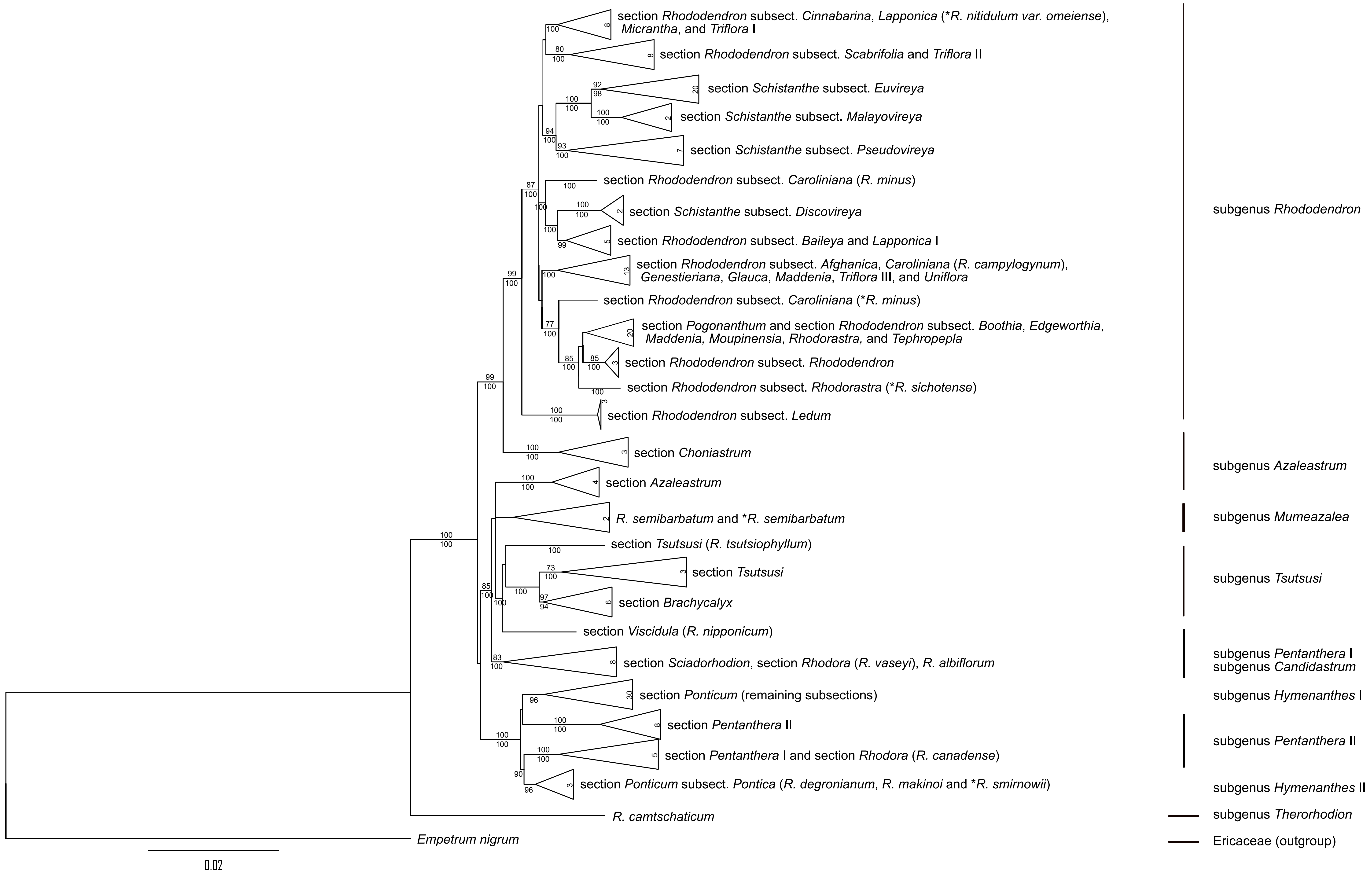

### Supplemental Figure 6

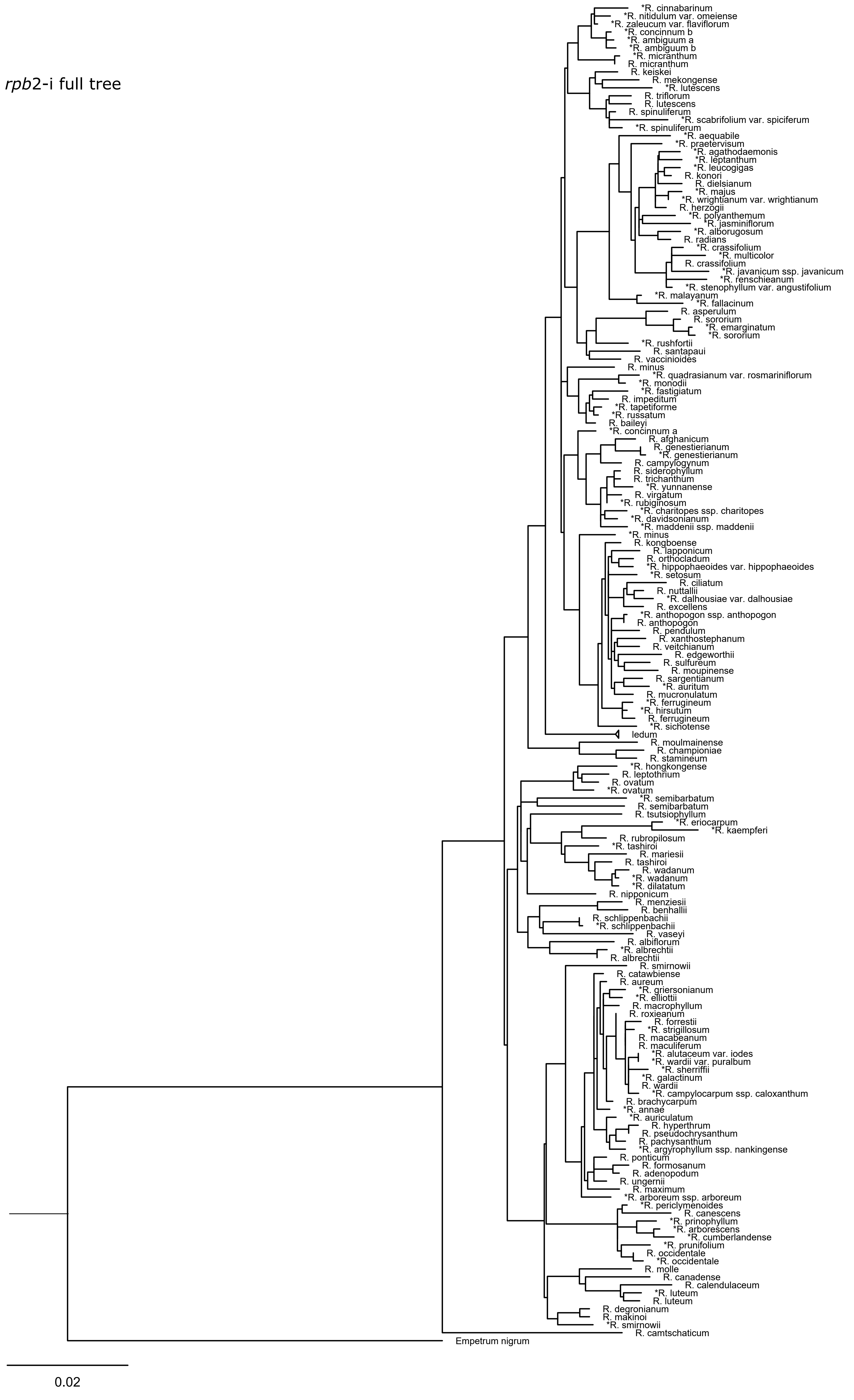

### Supplemental Figure 7

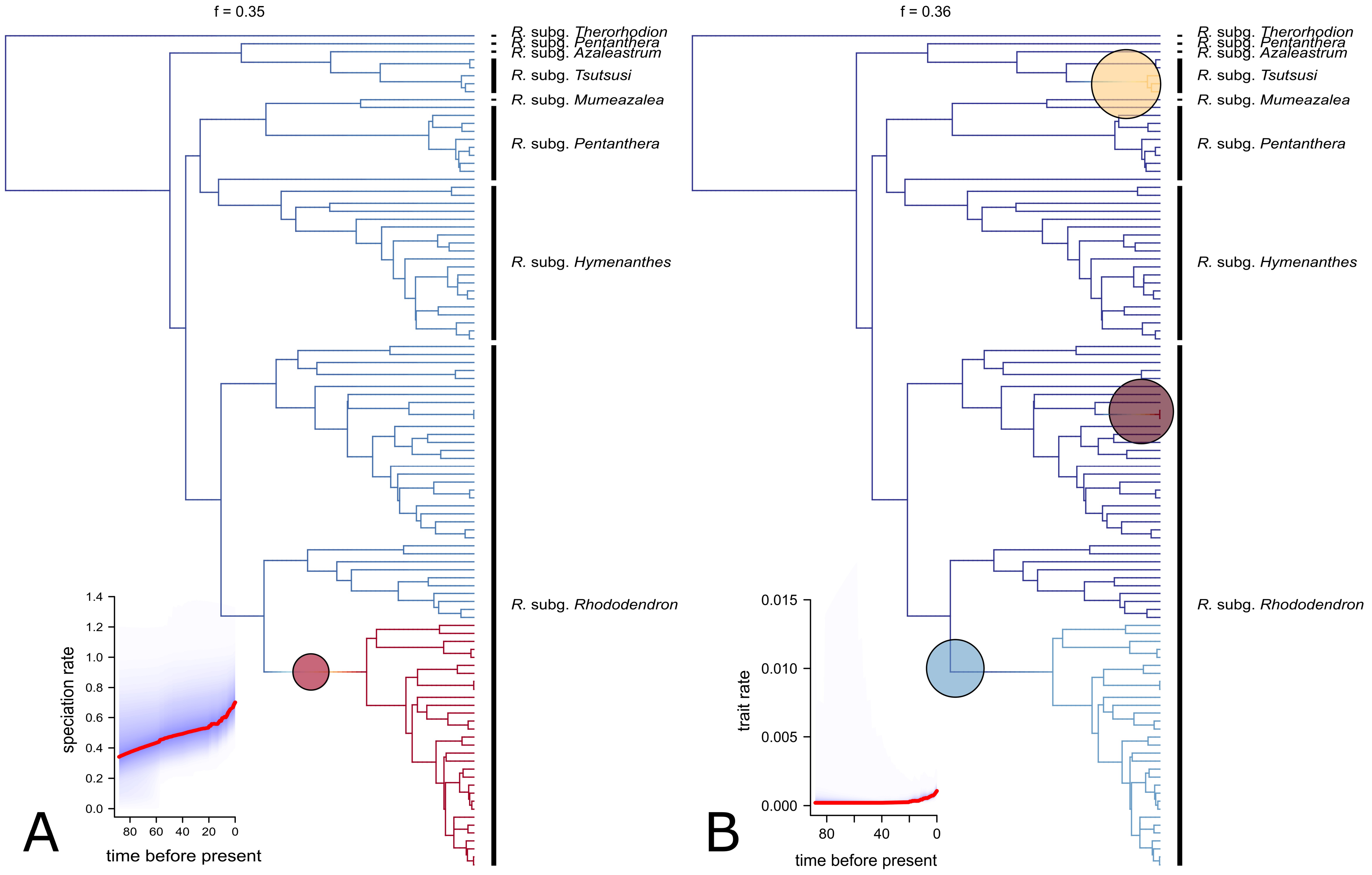

### Supplemental Figure 8

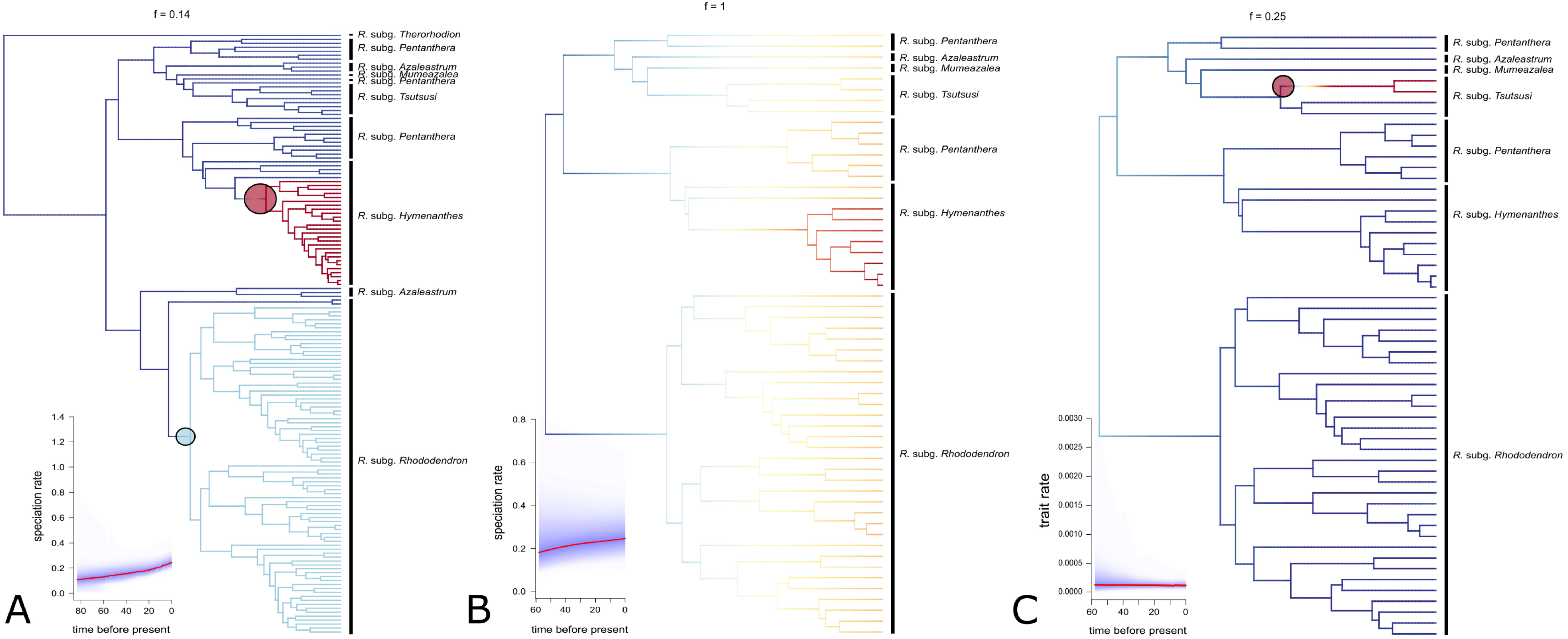

### Supplemental Figure 9

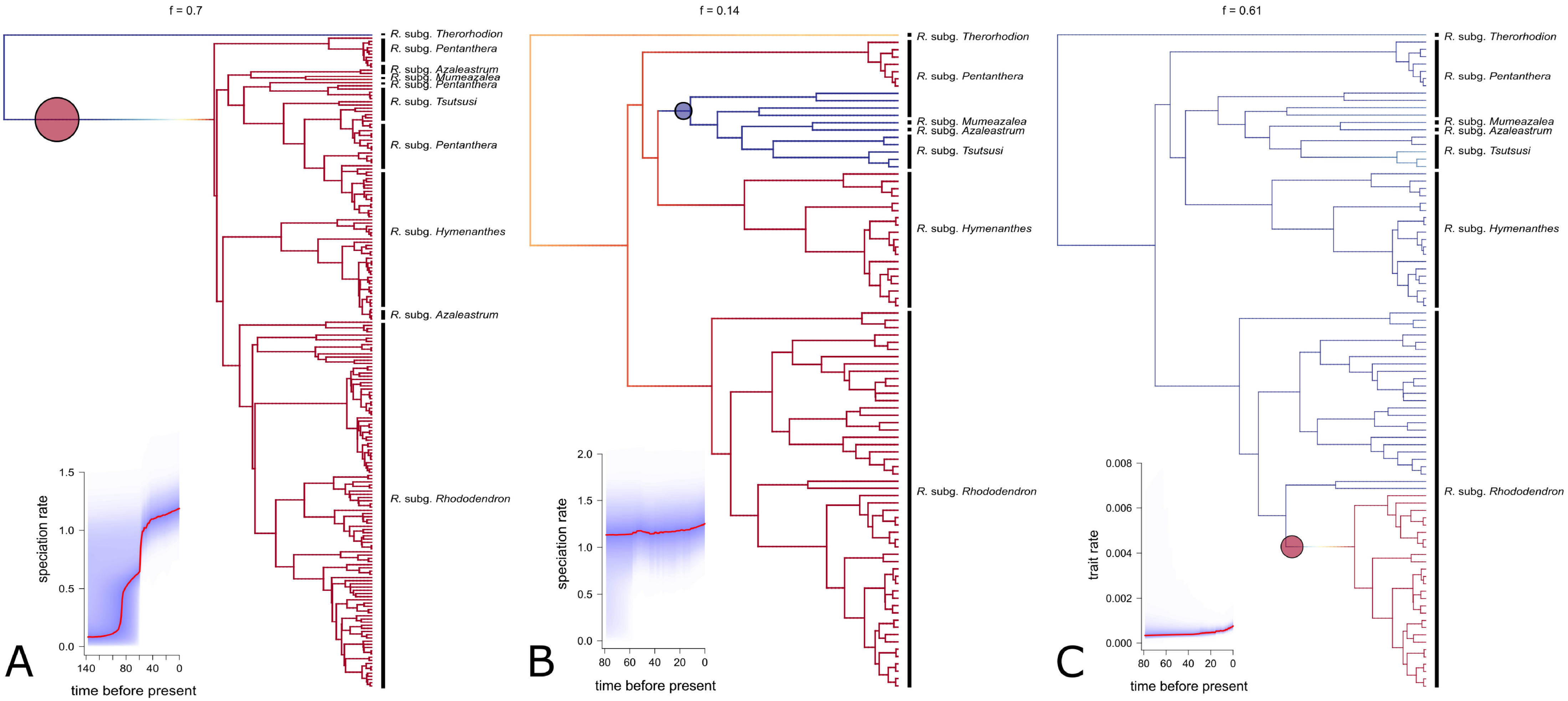

### Supplemental Figure 10

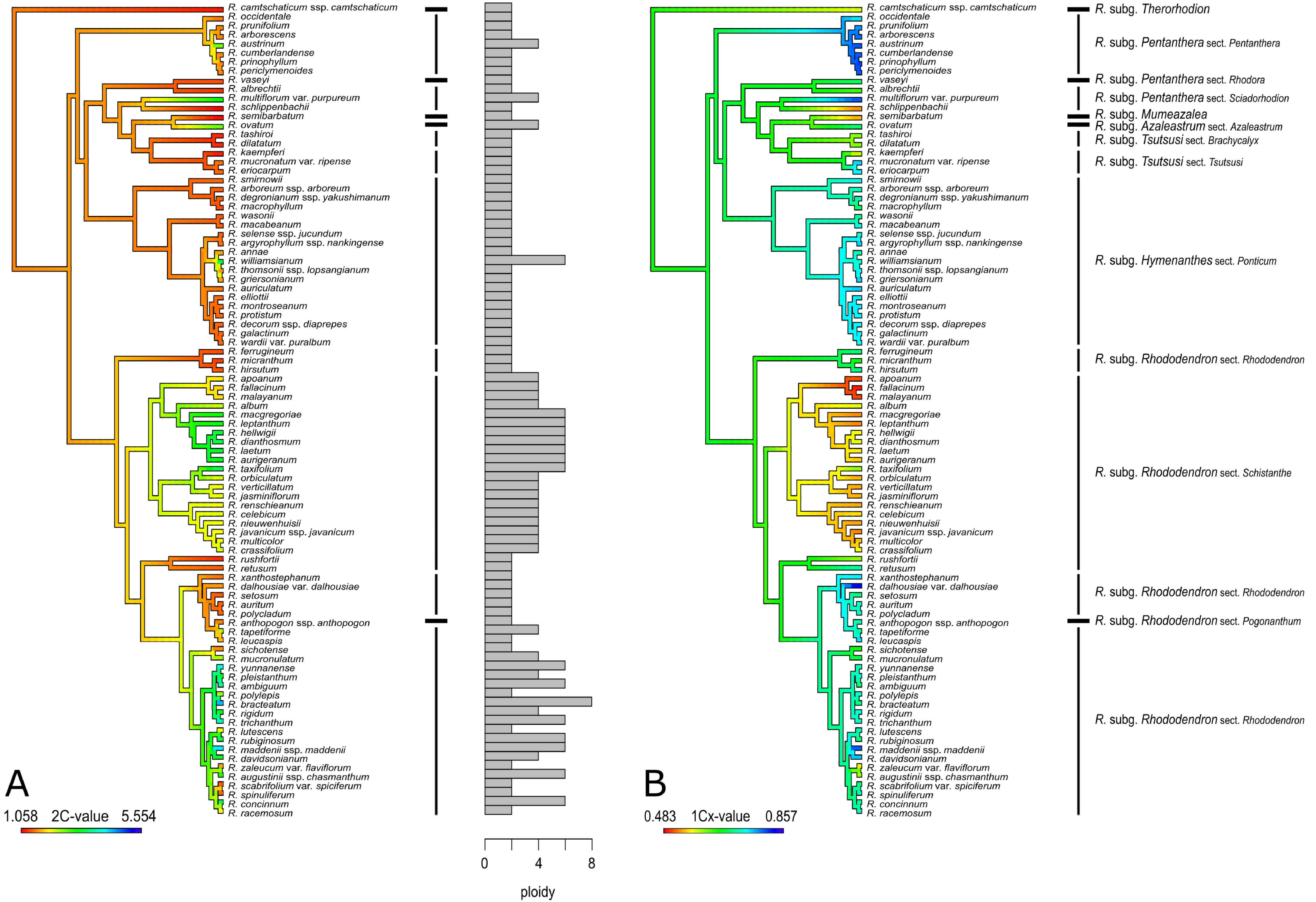
