## Supplemental Table 1 for "Incongruent phylogenies and its implications for the study of diversification, taxonomy and genome size evolution of *Rhododendron* (Ericaceae)"

| **TABLE S1.** *Rhododendron* subgenera and sections proposed by Chamberlain et al. (1996), Goetsch et al. (2005) and commented here. Species numbers are based on Chamberlain et al. (1996), which is, however, considered outdated nowadays partly since they used a wide species concept. Therefore, the species numbers were updated with information from Fang et al. (2005), Argent & Mitchell (2006) and checking the literature of species newly described in the genus since 1996 according to [www.ipni.org](http://www.ipni.org). *Diplarche*, *Ledum* and *Menziesia* have been included in the genus. This resulted in a preliminary checklist, which is available from the senior author on request but currently considered preliminary. | | | |
| --- | --- | --- | --- |
| **Chamberlain *et al.,* (1996)** | **Goetsch *et al.*, (2005)** | **Albach^(unpublished)^** | **No. species** |
| subgen. *Rhododendron* L. | Subgen. *Rhododendron* | Subgen. *Rhododendron* | 587 |
| sect. *Pogonanthum* Aitch. & Hemsl. |  |  |  |
| sect. *Rhododendron* L. |  |  |  |
| sect. *Schistanthe* Schltr. |  | sect. *Schistanthe* (excl. subsect. *Discovireya*) | (326) |
|  | incl. *Ledum* | excl. subsect. *Ledum* (unplaced to subgenus) | 8 |
| subgen. *Hymenanthes* (Blume) K. Koch | subgen. *Hymenanthes* | subgen. *Hymenanthes* |  |
| sect. *Ponticum* G. Don | sect. *Ponticum* | sect. *Ponticum* | 316 |
| subgen. *Pentanthera* (G. Don) Pojark. |  |  |  |
| sect. *Pentanthera* G. Don | sect. *Pentanthera*  (incl. sect. *Rhodora* p.p.) | subgen. *Pentanthera* s.str. (incl. sect. *Rhodora* p.p.) | 18 |
| sect. *Rhodora* (L.) G. Don |  | *R. vaseyi* (unplaced) | 1 |
|  | subgen. *Azaleastrum* | subgen. *Azaleastrum* | 154 |
| sect. *Sciadorhodion* Rehder & E.H. Wilson | sect. *Sciadorhodion* (incl. sect. *Rhodora* p.p., *Menziesia*) | sect. *Sciadorhodion* (incl. *Menziesia*, *R. schlippenbachii*) | (10) |
|  |  | other species (unplaced to subgenus) | 3 |
| sect. *Viscidula* Matsum. & Nakai | (incl. in sect. *Tsutsusi*) | *R. nipponicum* (unplaced to subgenus) | (1-) 2 |
| subgen. *Tsutsusi* (Sweet) Pojark. | sect. *Tsutsusi* | sect. *Tsutsusi* | (136) |
| sect. *Brachycaly x* Sweet |  |  |  |
| sect. *Tsutsusi* Sweet |  |  |  |
| subgen. *Azaleastrum* Planch. |  |  |  |
| sect. *Azaleastrum* (Planch.) Maxim. | (incl. in sect. *Tsutsusi*) | sect. *Azaleastrum* | (8) |
| sect. *Choniastrum* Franch. | subgen. *Choniastrum* | sect. *Choniastrum* (unplaced to subgenus) | 19 |
| subgen. *Candidastrum* Franch. | (incl. in sect. *Sciadorhodion*) | *R. albiflorum* (unplaced to subgenus) | 1 |
| subgen. *Mumeazalea* (Sleumer) W. R. Philipson & M.N.Philipson | (incl. in sect. *Tsutsusi*) | *R. semibarbatum* (unplaced to subgenus) | 1 |
| subgen. *Therorhodion* (Maxim.) A. Gray | subgen. *Therorhodion* (Maxim.) A. Gray | subgen. *Therorhodion* (Maxim.) A. Gray | 2 |
|  |  | Unplaced to subgenus (excl. mentioned above) | 44 |
|  |  | **Total no. of species** | **1156** |
