## Supplemental Table 2 for "Incongruent phylogenies and its implications for the study of diversification, taxonomy and genome size evolution of *Rhododendron* (Ericaceae)"

| **TABLE S2** Table showing information about species used in this work with accession numbers and flow cytometry results. *CV*: coefficient of variation; *ZM*: *Zea mays* (2.715pg/1C); *SP*: *Solanum pseudocapsicum* (1.29pg/1C); *HD*: *Hedychium gardnerianum* (2.01pg/1C); *Mbp*: mega base pairs | | | | | | | | | | | | | | | | |
| --- | --- | --- | --- | --- | --- | --- | --- | --- | --- | --- | --- | --- | --- | --- | --- | --- |
| **Subgenus** | **Section** | **Subsection** | **Series** | **Species** | **Herbarium and Accessions** | **Genbank Accession Number** | | | | | | **Genome Size** | | | | |
|  |  |  |  |  |  | ***trn*LF**^a^ | **nrITS**^b^ | ***mat*K-trnK intron**^c^ | ***rpb*2*i*-2** | ***rpb*2*i*-3** | ***rpb*2*i*-5** | **CV %** | **Standard** | **pg DNA** | **Ploidy Level** | **Mbp** |
| *Azaleastrum* | *Azaleastrum* |  |  | *honkongense* Hutchinson | 100470 | **✓** |  | **✓** |  | **✓** | **✓** |  |  |  |  |  |
|  |  |  |  | *leptothrium* Balf.f. & Forrest | WTU 357224 |  |  |  | AY765921 | AY765922 | AY765924 |  |  |  |  |  |
|  |  |  |  | *ovatum* (Lindley) Maximowicz | 100390 | **✓** | **✓** | **✓** |  | **✓** | **✓** | 5.91 | ZM | 2.812 | 4 | 2,755.760 |
|  |  |  |  | *ovatum* (Lindley) Maximowicz | NA |  |  | AB012729 |  |  |  |  |  |  |  |  |
|  |  |  |  | *ovatum* (Lindley) Maximowicz | WTU 357235 |  |  |  | AY765915 | AY765916 | AY765918 |  |  |  |  |  |
|  | *Choniastrum* |  |  | *championae* Hooker | 100863 | **✓** |  | **✓** |  |  |  | 3.08 | ZM | 1.165 | 2 | 1,141.373 |
|  |  |  |  | *championae* Hooker | WTU 357225 |  |  |  | AY765729 | AY765730 | AY765732 |  |  |  |  |  |
|  |  |  |  | *moulmainense* Hooker | 100886 | **✓** | **✓** | **✓** |  | **✓** |  |  |  |  |  |  |
|  |  |  |  | *moulmainense* Hooker | WTU 357234 |  |  |  | AY765723 | AY765724 | AY765726 |  |  |  |  |  |
|  |  |  |  | *stamineum* Franchet | WTU 357191 |  |  |  | AY765717 | AY765718 | AY765720 |  |  |  |  |  |
|  |  |  |  | *stamineum* Franchet | NA |  |  | AB012730 |  |  |  |  |  |  |  |  |
| *Candidastrum* |  |  |  | *albiflorum* Hooker | 2007/083d | **✓** | **✓** | **✓** |  |  |  |  |  |  |  |  |
|  |  |  |  | *albiflorum* Hooker | WTU 357195 |  |  |  | AY765975 | AY765976 | AY765978 |  |  |  |  |  |
|  |  |  |  | *albiflorum* Hooker |  |  |  | AB012731 |  |  |  |  |  |  |  |  |
| *Hymenanthes* | *Ponticum* | unplaced |  | *purdomii* Rehder & Wilson | 100796 |  |  | **✓** |  |  |  |  |  |  |  |  |
|  |  | *Arborea* |  | *arboreum* ssp. *arboreum* Smith | 100463 | **✓** | **✓** | **✓** | **✓** | **✓** |  | 3.76 | ZM | 1.459 | 2 | 1,429.493 |
|  |  |  |  | *arboreum* ssp. *delavayi* (Franchet) Chamberlain | 100466 |  |  | **✓** |  |  |  |  |  |  |  |  |
|  |  |  |  | *lanigerum* Tagg | RBGE 19291008 | EU087362 |  | EU087298 |  |  |  |  |  |  |  |  |
|  |  | *Argyrophylla* |  | *adenopodum* Franchet | RBGE 19620072 | EU087363 |  | EU087299 |  |  |  |  |  |  |  |  |
|  |  |  |  | *adenopodum* Franchet | WTU 357198 |  |  |  | AY765843 | AY765844 | AY765846 |  |  |  |  |  |
|  |  |  |  | *argyrophyllum* Franchet | RBGE 19810820 | EU087365 |  | EU087302 |  |  |  |  |  |  |  |  |
|  |  |  |  | *argyrophyllum* ssp. *nankingense* (Cowan) Chamberlain | 100812 | **✓** | **✓** | **✓** | **✓** | **✓** | **✓** | 5.45 | ZM | 1.557 | 2 | 1,525.533 |
|  |  |  |  | *coryanum* Tagg & Forrest | RBGE 19698489 | EU087364 |  | EU087300 |  |  |  |  |  |  |  |  |
|  |  |  |  | *formosanum* Hemsley | WTU 357214 |  |  |  | AY765849 | AY765850 | AY765852 |  |  |  |  |  |
|  |  |  |  | *insigne* Hemsley & Wilson | 100355 |  |  | **✓** |  |  |  |  |  |  |  |  |
|  |  |  |  | *insigne* Hemsley & Wilson | RBGE 19698662 | EU087301 |  | EU087301 |  |  |  |  |  |  |  |  |
|  |  |  |  | *ririei* Hemsley & Wilson | 100889 | **✓** | **✓** | **✓** |  |  |  | 4.67 | SP | 1.468 | 2 | 1,438.640 |
|  |  |  |  | *thayerianum* Rehder & Wilson | 100856 | **✓** | **✓** | **✓** |  |  |  | 5.23 | SP | 1.438 | 2 | 1,409.240 |
|  |  | *Auriculata* |  | *auriculatum* Hemsley | 100016 | **✓** | **✓** | **✓** | **✓** | **✓** | **✓** | 4.34 | ZM | 1.537 | 2 | 1,506.587 |
|  |  |  |  | *auriculatum* Hemsley | RBGE 19160027 | EU087366 |  | EU087303 |  |  |  | 3.77 | SP | 3.360 | 4 | 3,292.800 |
|  |  | *Barbata* |  | *barbatum* Don | RBGE 19370186 | EU087367 |  | EU087304 |  |  |  |  |  |  |  |  |
|  |  |  |  | *succothii* Davidian | RBGE 19902710 | EU087368 |  | EU087305 |  |  |  |  |  |  |  |  |
|  |  | *Campanulata* |  | *campanulatum* Don | OLD00685 |  |  | **✓** |  |  |  |  |  |  |  |  |
|  |  |  |  | *campanulatum* Don | RBGE 19720857 | EU087369 |  | EU087306 |  |  |  |  |  |  |  |  |
|  |  | *Campylocarpa* |  | *campylocarpum* Hooker | RBGE 19832543 | EU087370 |  | EU087307 |  |  |  |  |  |  |  |  |
|  |  |  |  | *campylocarpum* ssp. *caloxanthum* (Balfour & Farrer) Chamberlain | 100315 | **✓** | **✓** | **✓** | **✓** | **✓** | **✓** | 2.21 | SP | 1.408 | 2 | 1,379.840 |
|  |  |  |  | *wardii* Smith | RGBE 19698916 | EU087371 |  | EU087308 |  |  |  |  |  |  |  |  |
|  |  |  |  | *wardii* Smith | WTU 357175 |  |  |  | AY765735 | AY765736 | AY765738 |  |  |  |  |  |
|  |  |  |  | *wardii* var. *puralbum* (Balfour & Smith) Chamberlain | 100439 | **✓** | **✓** | **✓** | **✓** | **✓** |  | 2.41 | ZM | 1.487 | 2 | 1,456.933 |
|  |  | *Falconera* |  | *falconeri* Hooker | RBGE 19751301 | EU087372 |  | EU087309 |  |  |  |  |  |  |  |  |
|  |  |  |  | *falconeri* SPp. *eximium* (Nuttall) Chamberlain | 100473 |  | **✓** | S1 |  |  |  |  |  |  |  |  |
|  |  |  |  | *galactinum* Tagg | 100349 | **✓** | **✓** | **✓** | **✓** | **✓** |  | 4.76 | ZM | 1.511 | 2 | 1,480.429 |
|  |  |  |  | *galactinum* Tagg | RGBE 19913322 | EU087373 |  | EU087310 |  |  |  |  |  |  |  |  |
|  |  |  |  | *rothschildii* Davidian | RBGE 19764149 | EU087374 |  | EU087311 |  |  |  |  |  |  |  |  |
|  |  |  |  | *semnoides* Tagg & Forrest | RBGE 19301019 | EU087375 |  | EU087312 |  |  |  |  |  |  |  |  |
|  |  |  |  | *sinofalconeri* Balfour | RBGE 19960615 | EU087376 |  | EU087313 |  |  |  |  |  |  |  |  |
|  |  | *Fortunea* |  | *calophytum* Franchet | RBGE 19724038 | EU087379 |  | EU087316 |  |  |  |  |  |  |  |  |
|  |  |  |  | *decorum* Franchet | RBGE 19871529 | EU087380 |  | EU087317 |  |  |  |  |  |  |  |  |
|  |  |  |  | *decorum* SPp. *diaprepes* (Balfour & Smith) Ming | 100891 | **✓** | **✓** | **✓** | **✓** | **✓** |  | 3.59 | ZM | 1.506 | 2 | 1,475.880 |
|  |  |  |  | *orbiculare* Decaisne | RBGE 19460119 | EU087378 |  | EU087315 |  |  |  |  |  |  |  |  |
|  |  |  |  | *praevernum* Hutchinson | 100394 | **✓** | **✓** | **✓** | **✓** |  |  | 4.62 | SP | 1.521 | 2 | 1,490.580 |
|  |  |  |  | *praevernum* a Hutchinson | RBGE 1017*15376 | EU087381 |  | EU087318 |  |  |  |  |  |  |  |  |
|  |  |  |  | *praevernum* b Hutchinson | RBGE 19240357 | EU087382 |  | EU087319 |  |  |  |  |  |  |  |  |
|  |  |  |  | *vernicosum* Franchet | RBGE 19141012 | EU087383 |  | EU087320 |  |  |  |  |  |  |  |  |
|  |  | *Fulgensia* |  | *fulgens* Hooker | 100455 |  |  | **✓** |  |  |  |  |  |  |  |  |
|  |  |  |  | *fulgens* Hooker | RBGE 19371010 | EU087377 |  | EU087314 |  |  |  |  |  |  |  |  |
|  |  | *Fulva* |  | *fulvum* Balfour & Smith | RBGE 19180010 | EU087384 |  | EU087321 |  |  |  |  |  |  |  |  |
|  |  |  |  | *fulvum* SPp. *fulvum* | 100828 |  |  | **✓** |  |  |  |  |  |  |  |  |
|  |  | *Glischra* |  | *adenosum* Davidian | RBGE 19300433 | EU087389 |  | EU087326 |  |  |  |  |  |  |  |  |
|  |  |  |  | *crinigerum* Franchet | RBGE 19491014 | EU087390 |  | EU087327 |  |  |  |  |  |  |  |  |
|  |  |  |  | *glischrum* Balfour & Smith | RBGE 19950966 | EU087391 |  | EU087328 |  |  |  |  |  |  |  |  |
|  |  |  |  | *recurvoides* Tagg & Kingdon-Ward | 100840 |  |  | **✓** |  |  |  | 3.16 | ZM | 1.44 | 2 | 1,411.200 |
|  |  | *Grandia* |  | *balangense* Fang | 100018 |  |  | **✓** |  |  |  |  |  |  |  |  |
|  |  |  |  | *grande* Wight | RBGE 19698606 | EU087385 | GU176633 | EU087322 |  |  |  |  |  |  |  |  |
|  |  |  |  | *macabeanum* Balfour | 100467 | **✓** | **✓** | **✓** | **✓** |  |  | 4.26 | ZM | 1.507 | 2 | 1,476.533 |
|  |  |  |  | *macabeanum* Balfour | RBGE 19281023 | EU087386 |  | EU087323 |  |  |  |  |  |  |  |  |
|  |  |  |  | *macabeanum* Balfour | WTU 357238 |  |  |  | AY765741 | AY765742 | AY765744 |  |  |  |  |  |
|  |  |  |  | *montroseanum* Davidian | 100884 | **✓** | **✓** | **✓** |  |  |  | 4.34 | SP | 1.506 | 2 | 1,475.880 |
|  |  |  |  | *protistum* Balfour & Forrest | 100880 | **✓** | **✓** | **✓** |  |  |  | 2.98 | SP | 1.500 | 2 |  |
|  |  |  |  | *pudorosum* Cowan | RBGE 19764021 | EU087387 |  | EU087324 |  |  |  |  |  |  |  |  |
|  |  | *Griersoniana* |  | *griersonianum* Balfour & Forrest | 100483 | **✓** | **✓** | **✓** | **✓** | **✓** | **✓** | 3.75 | ZM | 1.558 | 2 | 1,527.106 |
|  |  |  |  | *griersonianum* Balfour & Forrest | RBGE 19320271 | EU087388 |  | EU087325 |  |  |  |  |  |  |  |  |
|  |  | *Irrorata* |  | *aberconwayi* Cowan | RBGE 19370338 | EU087392 |  | EU087329 |  |  |  |  |  |  |  |  |
|  |  |  |  | *annae* Franchet | 100794 | **✓** | **✓** | **✓** | **✓** | **✓** | **✓** | 3.08 | ZM | 1.462 | 2 | 1,432.760 |
|  |  |  |  | *irroratum* Franchet | RBGE 19812433 | EU087393 |  | EU087330 |  |  |  |  |  |  |  |  |
|  |  |  |  | *mengtszense* Balfour & Smith | RBGE 19960617 | EU087394 |  | EU087331 |  |  |  |  |  |  |  |  |
|  |  | *Lanata* |  | *lanatum* Hooker | RBGE 19810957 | EU087395 |  | EU087332 |  |  |  |  |  |  |  |  |
|  |  |  |  | *tsariense* Cowan | RBGE 19371016 | EU087396 |  | EU087333 |  |  |  |  |  |  |  |  |
|  |  | *Maculifera* |  | *anwheiense* Wilson | RBGE 19710038 | EU087397 |  | EU087334 |  |  |  |  |  |  |  |  |
|  |  |  |  | *maculiferum* Franchet | RBGE 19810812 | EU087398 |  | EU087335 |  |  |  |  |  |  |  |  |
|  |  |  |  | *maculiferum* Franchet | WTU 357164 |  |  |  | AY765777 | AY765778 | AY765780 |  |  |  |  |  |
|  |  |  |  | *pachysanthum* Hayata | WTU 357258 |  |  |  | AY765795 | AY765796 | AY765798 |  |  |  |  |  |
|  |  |  |  | *pseudochrysanthum* Hayata | RBGE 19810864 | EU087400 |  | EU087337 |  |  |  |  |  |  |  |  |
|  |  |  |  | *pseudochrysanthum* Hayata | WTU 357246 |  |  |  | AY765789 | AY765790 | AY765792 |  |  |  |  |  |
|  |  |  |  | *strigillosum* Franchet | 100420 | **✓** | **✓** | **✓** |  | **✓** | **✓** | 2.07 | SP | 1.486 | 2 | 1,456.280 |
|  |  |  |  | *strigillosum* Franchet | RBGE 19754050 | EU087399 |  | EU087336 |  |  |  |  |  |  |  |  |
|  |  | *Neriiflora* |  | *beanianum* Cowan | RBGE 19698404 | EU087401 |  | EU087338 |  |  |  |  |  |  |  |  |
|  |  |  |  | *catacosmum* Tagg | RBGE 19698450 | EU087402 |  | EU087339 |  |  |  |  |  |  |  |  |
|  |  |  |  | *citriniflorum* var. *citriniflorum* | 100323 |  |  | **✓** |  |  |  | 4.95 | ZM | 2.750 | 4 | 2,695.000 |
|  |  |  |  | *dichroanthum* Diels | RBGE 19200011 | EU087403 |  | EU087340 |  |  |  |  |  |  |  |  |
|  |  |  |  | *eudoxum* Forrest | RBGE 19794034 | EU087404 |  | EU087341 |  |  |  |  |  |  |  |  |
|  |  |  |  | *floccigerum* Franchet | RBGE 19491018 | EU087405 |  | EU087342 |  |  |  |  |  |  |  |  |
|  |  |  |  | *forrestii* Diels | RBGE 19250125 | EU087408 |  | EU087345 |  |  |  |  |  |  |  |  |
|  |  |  |  | *forrestii* Diels | WTU 357173 |  |  |  | AY765747 | AY765748 | AY765750 |  |  |  |  |  |
|  |  |  |  | *mallotum* Balfour & Kingdon-Ward | RBGE 19201013 | EU087406 |  | EU087343 |  |  |  |  |  |  |  |  |
|  |  |  |  | *microgynum* Balfour & Forrest | 100369 |  |  | **✓** |  |  |  |  |  |  |  |  |
|  |  |  |  | *neriiflorum* Franchet | 1999/259 | **✓** | **✓** | **✓** | **✓** |  |  | 4.40 | SP | 1.453 | 2 | 1,423.940 |
|  |  |  |  | *neriiflorum* Franchet | RBGE 19200019 | EU087407 |  | EU087344 |  |  |  |  |  |  |  |  |
|  |  | *Parishia* |  | *elliotii* Watt ex Brandis | 100496 | **✓** | **✓** | **✓** | **✓** | **✓** | **✓** | 4.04 | ZM | 1.514 | 2 | 1,483.720 |
|  |  | *Pontica* |  | *aureum* Georgi | 100015 |  |  | **✓** |  |  |  |  |  |  |  |  |
|  |  |  |  | *aureum* Georgi | RBGE 19450053 | AY496918 |  | AY494177 |  |  |  |  |  |  |  |  |
|  |  |  |  | *aureum* Georgi | WTU 357232 |  |  |  | AY765771 | AY765772 | AY765774 |  |  |  |  |  |
|  |  |  |  | *brachycarpum* Don | RBGE 19660135 | AY496917 |  | AY494176 |  |  |  |  |  |  |  |  |
|  |  |  |  | *brachycarpum* Don | WTU 357233 |  |  |  | AY765759 | AY765760 | AY765762 | 7.37 | SP | 1.363 | 2 | 1,335.740 |
|  |  |  |  | *brachycarpum* *SPp. brachycarpum* Don ex Don | 100248 | **✓** | **✓** | **✓** |  |  |  |  |  |  |  |  |
|  |  |  |  | *catawbiense* Michaux | RBGE 19340114 | AY496915 |  | AY494174 |  |  |  |  |  |  |  |  |
|  |  |  |  | *catawbiense* Michaux | WTU 357179 |  |  |  | AY765807 | AY765808 | AY765810 |  |  |  |  |  |
|  |  |  |  | *caucasicum* Pallas | RBGE 19521068 | AY496916 |  | AY494175 |  |  |  |  |  |  |  |  |
|  |  |  |  | *degronianum* spp. *degronianum* Carriére | RBGE 19341071 | AY496920 |  | AY494179 |  |  |  |  |  |  |  |  |
|  |  |  |  | *degronianum* spp. *degronianum* Carriére | WTU 357231 |  |  |  | AY765831 | AY765832 | AY765834 |  |  |  |  |  |
|  |  |  |  | *degronianum* SPp. *yakushimanum* (Nakai) Hara | 100338 | **✓** | **✓** | **✓** |  | **✓** | **✓** | 4.88 | SP | 1.479 | 2 | 1,449.271 |
|  |  |  |  | *hyperythrum* Hayata | RBGE 19410106 | AY496922 |  | AY494181 |  |  |  |  |  |  |  |  |
|  |  |  |  | *hyperythrum* Hayata | WTU 357217 |  |  |  | AY765801 | AY765802 | AY765804 |  |  |  |  |  |
|  |  |  |  | *macrophyllum* Don ex Don | 100837 | **✓** | **✓** | **✓** | **✓** |  |  | 2.86 | SP | 1.356 | 2 | 1,328.880 |
|  |  |  |  | *macrophyllum* Don ex Don | RBGE 19734184 | AY496914 |  | AY494173 |  |  |  |  |  |  |  |  |
|  |  |  |  | *macrophyllum* Don ex Don | WTU 351265 |  |  |  | AY765765 | AY765766 | AY765768 |  |  |  |  |  |
|  |  |  |  | *makinoi* Tagg | WTU 357180 |  |  |  | AY765837 | AY765838 | AY765840 |  |  |  |  |  |
|  |  |  |  | *maximum* Linneaus | RBGE 19800047 | AY496912 |  | AY494171 |  |  |  |  |  |  |  |  |
|  |  |  |  | *maximum* Linneaus | WTU 357194 |  |  |  | AY765819 | AY765820 | AY765822 |  |  |  |  |  |
|  |  |  |  | *ponticum* Linneaus | 100393 | **✓** | **✓** | **✓** |  | **✓** |  | 4.53 | SP | 1.452 | 2 | 1,422.960 |
|  |  |  |  | *ponticum* a Linneaus | RBGE 19773079 | AY496913 |  | AY494172 |  |  |  |  |  |  |  |  |
|  |  |  |  | *ponticum* b Linneaus | NA |  |  | AB012732 |  |  |  |  |  |  |  |  |
|  |  |  |  | *ponticum* Linneaus | WTU 357248 |  |  |  | AY765813 | AY765814 | AY765816 |  |  |  |  |  |
|  |  |  |  | *smirnowii* Trautvetter | 100417 | **✓** | **✓** | **✓** | **✓** | **✓** | **✓** | 2.94 | ZM | 1.487 | 2 | 1,456.933 |
|  |  |  |  | *smirnowii* Trautvetter | RBGE 19698845 | AY496921 |  | AY494180 |  |  |  |  |  |  |  |  |
|  |  |  |  | *smirnowii* Trautvetter | WTU 357196 |  |  |  | AY765825 | AY765826 | AY765828 |  |  |  |  |  |
|  |  |  |  | *ungernii* Trautvetter | RBGE 19623836 | AY496919 |  | AY494178 |  |  |  |  |  |  |  |  |
|  |  |  |  | *ungernii* Trautvetter | WTU 357183 |  |  |  | AY765783 | AY765784 | AY765786 |  |  |  |  |  |
|  |  | *Selensia* |  | *selense* SPp. *jucundum* (Balfour & Smith) Chamberlain | 100412 | **✓** | **✓** | **✓** |  | **✓** |  | 2.27 | ZM | 1.437 | 2 | 1,408.587 |
|  |  |  |  | *selense* Franchet | RBGE 19812509 | EU087409 |  | EU087346 |  |  |  |  |  |  |  |  |
|  |  | *Taliensia* |  | *aganniphum* Balfour & Kingdon-Ward | RBGE 20052580 | EU087412 |  | EU087349 |  |  |  |  |  |  |  |  |
|  |  |  |  | *aganniphum* var. *flavorufum* (Balfour & Forrest) Chamberlain | 100830 |  |  | **✓** |  |  |  |  |  |  |  |  |
|  |  |  |  | *alutaceum* Balfour & Smith | RBGE 19913281 | EU087410 |  | EU087347 |  |  |  |  |  |  |  |  |
|  |  |  |  | *alutaceum* var. *iodes* (Balfour & Forrest) Chamberlain | 100821 | **✓** | **✓** | **✓** | **✓** |  | **✓** | 6.00 | SP | 1.463 | 2 | 1,433.740 |
|  |  |  |  | *beesianum* Diels | 100824 |  |  | **✓** |  |  |  |  |  |  |  |  |
|  |  |  |  | *beesianum* Diels | RBGE 19491013 | EU087411 |  | EU087348 |  |  |  |  |  |  |  |  |
|  |  |  |  | *bureavii* Franchet | RBGE 19331022 | EU087416 |  | EU087353 |  |  |  |  |  |  |  |  |
|  |  |  |  | *faberi* Hemsley | 101066 |  |  | **✓** |  |  |  |  |  |  |  |  |
|  |  |  |  | *nakotiltum* Balfour & Forrest | RBGE 1793*28170 | EU087413 |  | EU087350 |  |  |  |  |  |  |  |  |
|  |  |  |  | *phaeochrysum* Balfour & Smith | RBGE 19698781 | EU087414 |  | EU087351 |  |  |  |  |  |  |  |  |
|  |  |  |  | *roxieanum* Forrest | RBGE 19734059 | EU087417 |  | EU087354 |  |  |  |  |  |  |  |  |
|  |  |  |  | *roxieanum* Forrest | WTU 357228 |  |  |  | AY765753 | AY765754 | AY765756 |  |  |  |  |  |
|  |  |  |  | *taliense* Franchet | RBGE 19568652 | EU087415 |  | EU087352 |  |  |  |  |  |  |  |  |
|  |  |  |  | *wasonii* a Hemsley & Wilson | 100799 | **✓** | **✓** | **✓** |  |  |  | 2.68 | SP | 1.43 | 2 | 1,401.400 |
|  |  |  |  | *wasonii* b Hemsley & Wilson | 100860 | **✓** | **✓** | **✓** |  |  |  |  |  |  |  |  |
|  |  | *Thomsonia* |  | *cerasinum* Tagg | RBGE 19291004 | EU087418 |  | EU087355 |  |  |  |  |  |  |  |  |
|  |  |  |  | *eclecteum* Balfour & Forrest | RBGE 19201020 | EU087419 |  | EU087356 |  |  |  |  |  |  |  |  |
|  |  |  |  | *faucium* Chamberlain | RBGE 19470106 | EU087420 |  | EU087357 |  |  |  |  |  |  |  |  |
|  |  |  |  | *hookeri* Nuttall | RBGE 19291007 | EU087421 |  | EU087358 |  |  |  |  |  |  |  |  |
|  |  |  |  | *hylaeum* Balfour & Farrer | 100489 | **✓** | **✓** | **✓** |  |  |  | 1.98 | SP | 1.425 | 2 | 1,396.500 |
|  |  |  |  | *sheriffii* Cowan | 100795 | **✓** | **✓** | **✓** | **✓** | **✓** | **✓** | 5.53 | SP | 1.438 | 2 | 1,409.240 |
|  |  |  |  | *subansiriense* Chamberlain | 100877 |  |  | **✓** |  |  |  |  |  |  |  |  |
|  |  |  |  | *thomsonii* Hooker | RBGE 19803353 | EU087422 |  | EU087359 |  |  |  |  |  |  |  |  |
|  |  |  |  | *thomsonii* SPp. *lopsangianum* Hooker | 100491 | **✓** | **✓** | **✓** | **✓** |  |  | 4.34 | ZM | 1.479 | 2 | 1,449.582 |
|  |  | *Venatora* |  | *venator* Tagg | 2010/238 |  |  | **✓** |  |  |  | 3.32 | ZM | 1.434 | 2 | 1,405.320 |
|  |  |  |  | *venator* Tagg | RBGE 19370170 | EU087423 |  | EU087360 |  |  |  |  |  |  |  |  |
|  |  | *Williamsiana* |  | *williamsianum* Rehder & Wilson | 100441 |  | **✓** | **✓** |  |  |  | 3.00 | ZM | 2.724 | 4 | 2,669.520 |
|  |  |  |  | *williamsianum* Rehder & Wilson | RBGE 19320138 | EU087424 |  | EU087361 |  |  |  |  |  |  |  |  |
| *Mumeazalea* |  |  |  | *semibarbatum* Maximowicz | 100781 | **✓** | **✓** | **✓** | **✓** | **✓** | **✓** | 3.06 | ZM | 1.079 | 2 | 1,057.093 |
|  |  |  |  | *semibarbatum* Maximowicz | NA |  |  | AB012733 |  |  |  |  |  |  |  |  |
|  |  |  |  | *semibarbatum* Maximowicz | WTU 357216 |  |  |  | AY765933 | AY765934 | AY765936 |  |  |  |  |  |
| *Pentanthera* | *Pentanthera* |  |  | *arborescens* (Pursh) Torrey | 101044 | **✓** | **✓** | **✓** | **✓** | **✓** |  | 3.38 | SP | 1.611 | 2 | 1,578.780 |
|  |  |  |  | *atlanticum* a (Ashe) Rehder | 100012 | **✓** | **✓** | **✓** |  |  |  | 3.56 | ZM | 3.828 | 4 | 3,751.440 |
|  |  |  |  | *atlanticum* b (Ashe) Rehder | 100013 | **✓** |  | **✓** |  |  |  |  |  |  |  |  |
|  |  |  |  | *atlanticum* (Ashe) Rehder | RBGE 19730782 | AY496924 |  | AY494183 |  |  |  |  |  |  |  |  |
|  |  |  |  | *austrinum (*Small) Rehder | 100456 | **✓** | **✓** | **✓** |  |  |  | 4.37 | SP | 3.182 | 4 | 3,118.948 |
|  |  |  |  | *calendulaceum* (Michaux) Torrey |  |  | GU176632 |  |  |  |  |  |  |  |  |  |
|  |  |  |  | *calendulaceum* (Michaux) Torrey | RSF SEH-1016 |  |  |  | AY765867 | AY765868 | AY765870 |  |  |  |  |  |
|  |  |  |  | *canescens* (Michaux) Sweet | WTU 357189 |  |  |  | AY765855 | AY765856 | AY765858 | 5.43 | HD | 1.662 | 2 | 1,628.760 |
|  |  |  |  | *cumberlandense* Braun | 101045 | **✓** | **✓** | **✓** | **✓** | **✓** |  | 3.58 | SP | 1.608 | 2 | 1,575.840 |
|  |  |  |  | *luteum* Sweet | 100363 | **✓** | **✓** | **✓** | **✓** |  | **✓** | 4.93 | ZM | 2.760 | 4 | 2,704.800 |
|  |  |  |  | *luteum* Sweet | RBGE 19773072 | AY496923 |  | AY494182 |  |  |  |  |  |  |  |  |
|  |  |  |  | *luteum* Sweet | WTU 357241 |  |  |  | AY765873 | AY765874 | AY765876 |  |  |  |  |  |
|  |  |  |  | *molle* (Blume) Don | WTU 357229 |  |  |  | AY765879 | AY765880 | AY765882 |  |  |  |  |  |
|  |  |  |  | *occidentale* (Torrey & Gray) Gray | 100380 | **✓** | **✓** | **✓** | **✓** | **✓** |  | 2.43 | SP | 1.50 | 2 | 1,470.000 |
|  |  |  |  | *occidentale* (Torrey & Gray) Gray | RBGE 19582084 | AY496925 |  | AY494184 |  |  |  |  |  |  |  |  |
|  |  |  |  | *occidentale* (Torrey & Gray) Gray | WTU 357230 |  |  |  | AY765957 | AY765958 | AY765960 |  |  |  |  |  |
|  |  |  |  | *periclymenoides* (Michaux) Shinners | 100391 | **✓** | **✓** | **✓** |  | **✓** | **✓** | 3.16 | SP | 1.611 | 2 | 1,579.270 |
|  |  |  |  | *prinophyllum* (Small) Millais | 100397 | **✓** | **✓** | **✓** | **✓** | **✓** |  | 5.09 | ZM | 1.646 | 2 | 1,613.080 |
|  |  |  |  | *prunifolium* (Small) Millais | 101046 | **✓** | **✓** | **✓** | **✓** | **✓** |  | 3.53 | SP | 1.577 | 2 | 1,545,460 |
|  |  |  |  | *viscosum* (Linneaus) Torrey | 100434 |  |  | **✓** |  |  |  | 4.55 | ZM | 1.650 | 2 | 1,617.000 |
|  | *Rhodora* |  |  | *canadense* (Linneaus) Torrey | 101115 | **✓** |  |  |  |  |  | 7.21 | ZM | 1.202 | 2 | 1,177.960 |
|  |  |  |  | *canadense* (Linneaus) Torrey | NA |  |  | AB012735 |  |  |  |  |  |  |  |  |
|  |  |  |  | *canadense* (Linneaus) Torrey | WTU 357205 |  |  |  | AY765885 | AY765886 | AY765888 |  |  |  |  |  |
|  |  |  |  | *vaseyi* Gray | 100431 | **✓** | **✓** | **✓** |  | **✓** |  | 3.38 | SP | 1.392 | 2 | 1,364.160 |
|  |  |  |  | *vaseyi* Gray | WTU 357188 |  |  |  | AY765957 | AY765958 | AY765960 |  |  |  |  |  |
|  | *Sciadorhodion* |  |  | *albrechtii* Maximowicz | 100005 | **✓** | **✓** | **✓** | **✓** | **✓** | **✓** | 5.47 | SP | 1.291 | 2 | 1,265.180 |
|  |  |  |  | *albrechtii* Maximowicz | NA |  |  | AB012737 |  |  |  |  |  |  |  |  |
|  |  |  |  | *albrechtii* Maximowicz | WTU 357199 |  |  |  | AY765963 | AY765964 | AY765966 |  |  |  |  |  |
|  |  |  |  | *benhallii* Craven, *nom.nov.* (*Menziesia ciliicalyx* (Miquel) Maximowicz) | E 1969-5350 |  | EU855846 |  |  |  |  |  |  |  |  |  |
|  |  |  |  | *benhallii* Craven, *nom.nov.* (*Menziesia ciliicalyx* (Miquel) Maximowicz) | WTU 357162 |  |  |  | AY765951 | AY765952 | AY765954 |  |  |  |  |  |
|  |  |  |  | *benhallii* Craven var. *purpurea* (*Menziesia ciliicalyx* var. *purpurea*) | OLD00747 | **✓** | **✓** |  |  |  |  | 7.70 | ZM | 3.250 | 4 | 3,185.000 |
|  |  |  |  | *menziesii* Craven, *nom. nov.* (Menziesia *ferruginea* Smith) | UTW - Olmstead |  | EU855847 |  |  |  |  |  |  |  |  |  |
|  |  |  |  | *menziesii* Craven, *nom. nov.* (Menziesia *ferruginea* Smith) | WTU 357209 |  |  |  | AY765945 | AY765946 | AY765948 |  |  |  |  |  |
|  |  |  |  | *Menziesia lasiophylla* Nakai | RSF - s.n. |  | EU855848 |  |  |  |  |  |  |  |  |  |
|  |  |  |  | *nipponicum* Matsumura | NA |  |  | AB012739 |  |  |  |  |  |  |  |  |
|  |  |  |  | *nipponicum* Matsumura | WTU 357244 |  |  |  | AY765939 | AY765940 | AY765942 |  |  |  |  |  |
|  |  |  |  | *pentaphyllum* Maximowicz | NA |  |  | AB012738 |  |  |  |  |  |  |  |  |
|  |  |  |  | *pilosum* Craven, *nom.nov.* (*Menziesia pilosa* (Michaux) JuSPieu) | 1266 | **✓** | **✓** | **✓** | **✓** |  |  | 3.85 | ZM | 2.75 | 4 | 2,695.000 |
|  |  |  |  | *pilosum* Craven, *nom.nov.* (*Menziesia pilosa* (Michaux) JuSPieu) | WFU - Thornton |  | EU855849 |  |  |  |  |  |  |  |  |  |
|  |  |  |  | *kroniae* Craven, *nom.nov.* (*Menziesia purpurea* Maximowicz) | E - s.n. |  | EU855850 |  |  |  |  |  |  |  |  |  |
|  |  |  |  | *quinquefolium* BiSPet & Moore | 2010/213 |  |  | **✓** |  |  |  |  |  |  |  |  |
|  |  |  |  | *schlippenbachii* Maximowicz | 100408 | **✓** | **✓** | **✓** | **✓** | **✓** | **✓** | 3.74 | SP | 1.087 | 2 | 1,065.260 |
|  |  |  |  | *schlippenbachii* Maximowicz | NA |  |  | AB012736 |  |  |  |  |  |  |  |  |
| *Rhododendron* | *Pogonanthum* |  |  | *anthopogon* Don | WTU 357240 |  |  |  | AY765543 | AY765544 | AY765546 | 4.95 | HD | 1.404 | 2 | 1,375.920 |
|  |  |  |  | *anthopogon* Don SPp. *anthopogon* | 2006/232 | **✓** | **✓** | **✓** | **✓** | **✓** | **✓** |  |  |  |  |  |
|  |  |  |  | *kongboense* Hutchinson | WTU 357259 |  |  |  | AY765537 | AY765538 | AY765540 |  |  |  |  |  |
|  |  |  |  | *primuliflorum* Bureau & Franchet | NA |  |  | AB012740 |  |  |  |  |  |  |  |  |
|  |  |  |  | *sargentianum* Rehder & Wilson | WTU 357247 |  |  |  | AY765531 | AY765532 | AY765534 |  |  |  |  |  |
|  | *Rhododendron* |  |  | *afghanicum* Aitch. & Hemsley | WTU 357227 |  |  |  | AY765657 | AY765658 | AY765660 |  |  |  |  |  |
|  |  | *Baileya* |  | *baileyi* Balfour | 100017 |  |  | **✓** |  |  |  | 3.64 | ZM | 1.378 | 2 | 1,350.440 |
|  |  |  |  | *baileyi* Balfour | WTU 357181 |  |  |  | AY765573 | AY765574 | AY765576 |  |  |  |  |  |
|  |  | *Boothia* |  | *leucaspis* Tagg | 100468 | **✓** | **✓** | **✓** |  |  |  | 4.05 | ZM | 1.573 | 2 | 1,541.867 |
|  |  |  |  | *sulfureum* Franchet | WTU 357172 |  |  |  | AY765489 | AY765490 | AY765492 |  |  |  |  |  |
|  |  |  |  | *campylognum* Franchet | WTU 357215 |  |  |  | AY765633 | AY765634 | AY765636 |  |  |  |  |  |
|  |  | *Caroliniana* |  | *minus* Michaux | 100370 | **✓** |  | **✓** | **✓** | **✓** | **✓** | 2.79 | SP | 1.365 | 2 | 1,337.700 |
|  |  |  |  | *minus* var. *chapmanii* (Gray) Duncan & Pullen | WTU 357201 |  |  |  | AY765561 | AY765562 | AY765564 |  |  |  |  |  |
|  |  | *Cinnabarina* |  | *cinnabarinum* Hooker | 100322 | **✓** | **✓** | **✓** | **✓** | **✓** | **✓** | 1.98 | PS | 7.05 | 6 | 6,909.000 |
|  |  |  |  | *keysii* Nuttall | 100748 | **✓** | **✓** | **✓** |  | **✓** |  |  |  |  |  |  |
|  |  | *Edgeworthia* |  | *edgeworthii* Hooker | WTU 357186 |  |  |  | AY765483 | AY765484 | AY765486 |  |  |  |  |  |
|  |  |  |  | *pendulum* Hooker | 100487 |  |  | **✓** |  |  |  | 4.29 | ZM | 1.670 | 2 | 1,636.600 |
|  |  |  |  | *pendulum* Hooker | WTU 357203 |  |  |  | AY765495 | AY765496 | AY765498 |  |  |  |  |  |
|  |  | *Fragariflora* |  | *fragariflorum* Kingdon-Ward | 100908 |  |  | **✓** |  |  |  |  |  |  |  |  |
|  |  | *Genestieriana* |  | *genestierianum* Forrest | 100484 | **✓** | **✓** | **✓** |  | **✓** | **✓** | 5.50 | SP | 1.460 | 2 | 1,430.800 |
|  |  |  |  | *genestierianum* Forrest | WTU 357208 |  |  |  | AY765639 | AY765640 | AY765642 |  |  |  |  |  |
|  |  | *Glauca* |  | *charitopes* SPp. *charitopes* | 100866 | **✓** | **✓** | **✓** | **✓** | **✓** | **✓** |  |  |  |  |  |
|  |  |  |  | *charitopes* SPp. *tsangpoense* | 2001/1057 |  | **✓** | **✓** |  |  |  | 3.86 | HD | 1.324 | 2 | 1,297.520 |
|  |  |  |  | *luteiflorum* (Davidian) Cullen | 101051 |  |  | **✓** |  |  |  |  |  |  |  |  |
|  |  | *Heliolepida* |  | *bracteatum* Rehder & Wilson | 100848 | **✓** | **✓** | **✓** |  | **✓** |  | 3.17 | HD | 5.436 | 6 | 5,327.280 |
|  |  |  |  | *heliolepis* var. *fumidum* (Balfour & Smith) Fang | 2008/111 |  |  | **✓** |  |  |  | 2.08 | ZM | 4.254 | 6 | 4,168.920 |
|  |  |  |  | *rubiginosum* Franchet | 100404 | **✓** | **✓** | **✓** | **✓** | **✓** |  | 4.58 | ZM | 4.186 | 6 | 4,102.717 |
|  |  | *Lapponica* |  | *cuneatum* Smith | 100328 |  |  | **✓** |  |  |  | 3.41 | ZM | 4.020 | 6 | 3,939.600 |
|  |  |  |  | *fastigiatum* Franchet | 100343 | **✓** | **✓** | **✓** | **✓** | **✓** | **✓** |  |  |  |  |  |
|  |  |  |  | *hippophaeoides* Balfour & Smith | RBGE 19321022 |  | GU176634 |  |  |  |  |  |  |  |  |  |
|  |  |  |  | *hippophaeoides* var. *hippophaeoides* Hutchinson | 100353 | **✓** | **✓** | **✓** | **✓** | **✓** |  |  |  |  |  |  |
|  |  |  |  | *impeditum* Balfour & Smith | WTU 357219 |  |  |  | AY765567 | AY765568 | AY765570 |  |  |  |  |  |
|  |  |  |  | *lapponicum* (Linneaus) Wahlenberg | WTU 357176 |  |  |  | AY765519 | AY765520 | AY765522 |  |  |  |  |  |
|  |  |  |  | *nitidulum* var. *omeiense* Philipson & Philipson | 100377 | **✓** | **✓** | **✓** | **✓** | **✓** | **✓** |  |  |  |  |  |
|  |  |  |  | *orthocladum* Balfour & Forrest | WTU 357207 |  |  |  | AY765525 | AY765526 | AY765528 |  |  |  |  |  |
|  |  |  |  | *polycladum* Franchet | 100392 | **✓** | **✓** | **✓** |  | **✓** |  |  |  | 1.467 | 2 | 1,437.987 |
|  |  |  |  | *ruSPatum* Balfour & Forrest | 100384 | **✓** | **✓** | **✓** |  | **✓** | **✓** |  |  |  |  |  |
|  |  |  |  | *setosum* Don | 2010/384 | **✓** | **✓** | **✓** | **✓** | **✓** |  | 4.53 | ZM | 1.386 | 2 | 1,358.280 |
|  |  |  |  | *tapetiforme* Balfour & Kingdon-Ward | 101054 | **✓** | **✓** | **✓** | **✓** | **✓** |  | 3.56 | ZM | 2.749 | 4 | 2,694.347 |
|  |  | *Ledum* |  | *diversipilosum* (Nakai) Harmaja | NA |  |  | AB012751 |  |  |  |  |  |  |  |  |
|  |  |  |  | *hypoleucum* (Komarov) Harmaja | WTU 357226 |  |  |  | AY765705 | AY765706 | AY765708 |  |  |  |  |  |
|  |  |  |  | *tomentosum* (Stokes) Harmaja | 100426 | **✓** | **✓** | **✓** | **✓** | **✓** | **✓** | 2.83 | ZM | 2.442 | 4 | 2,393.160 |
|  |  |  |  | *tomentosum* (Stokes) Harmaja | WTU 357165 |  |  |  | AY765711 | AY765712 | AY765714 |  |  |  |  |  |
|  |  | *Lepidota* |  | *lepidotum* Wallich ex Don | 092b |  |  | **✓** |  |  |  |  |  |  |  |  |
|  |  | *Maddenia* |  | *carneum* Hutchinson | 100475 |  |  | **✓** |  |  |  | 3.77 | ZM | 2.766 | 4 | 2,710.680 |
|  |  |  |  | *ciliatum* Hooker | WTU 357237 |  |  |  | AY765513 | AY765514 | AY765516 |  |  |  |  |  |
|  |  |  |  | *dalhousiae* Hooker var. *dalhousiae* | 100486 | **✓** | **✓** | **✓** | **✓** | **✓** |  | 2.13 | ZM | 1.714 | 2 | 1,679.720 |
|  |  |  |  | *excellens* Hooker | WTU 357236 |  |  |  | AY765507 | AY765508 | AY765510 |  |  |  |  |  |
|  |  |  |  | *lindleyi* Moore | 100751 |  |  | **✓** |  |  |  | 3.22 | ZM | 1.682 | 2 | 1,648.360 |
|  |  |  |  | *maddenii* SPp. *maddenii* Hooker | 100750 | **✓** | **✓** | **✓** | **✓** | **✓** |  | 3.38 | SP | 5.056 | 6 | 4,954.880 |
|  |  |  |  | *nuttallii* Booth | WTU 357218 |  |  |  | AY765501 | AY765502 | AY765504 |  |  |  |  |  |
|  |  |  |  | *scopulorum* Hutchinson | 2002/1084 |  |  | **✓** |  |  |  |  |  |  |  |  |
|  |  |  |  | *veitchianum* Hooker | WTU 357202 |  |  |  | AY765477 | AY765478 | AY765480 |  |  |  |  |  |
|  |  | *Micrantha* |  | *micranthum* Turczaninow | 100368 | **✓** | **✓** | **✓** | **✓** |  | **✓** | 2.88 | ZM | 1.267 | 2 | 1,241.333 |
|  |  |  |  | *micranthum* Turczaninow | WTU 357260 |  |  |  | AY772486 | AY772487 | AY772489 |  |  |  |  |  |
|  |  | *Monantha* |  | *monanthum* Balfour & Smith | 2002/1079 |  |  | **✓** |  |  |  |  |  |  |  |  |
|  |  | *Moupinensia* |  | *moupinense* Franchet | 100374 | **✓** | **✓** | **✓** |  |  |  | 3.69 | ZM | 1.630 | 2 | 1,597.400 |
|  |  |  |  | *moupinense* Franchet | WTU 357243 |  |  |  | AY765465 | AY765466 | AY765468 |  |  |  |  |  |
|  |  | *Rhododendron* |  | *ferrugineum* Linneaus | 100345 | **✓** | **✓** | **✓** | **✓** | **✓** | **✓** | 4.16 | ZM | 1.407 | 2 | 1,379.293 |
|  |  |  |  | *ferrugineum* Linneaus | NA |  |  | AB012741 |  |  |  |  |  |  |  |  |
|  |  |  |  | *ferrugineum* Linneaus | WTU 357174 |  |  |  | AY765555 | AY765556 | AY765558 |  |  |  |  |  |
|  |  |  |  | *hirsutum* Linneaus | OLD00789 | **✓** | **✓** | **✓** | **✓** | **✓** |  | 3.40 | ZM | 1.462 | 2 | 1,432.760 |
|  |  | *Rhodorastra* |  | *ledebourii* Pojarkova | 100882 | **✓** | **✓** | **✓** |  |  |  |  |  |  |  |  |
|  |  |  |  | *mucronulatum* Turczaninow | 100376 | **✓** | **✓** | **✓** |  |  |  | 3.95 | ZM | 2.748 | 4 | 2,693.040 |
|  |  |  |  | *mucronulatum* Turczaninow | WTU 357184 |  |  |  | AY765549 | AY765550 | AY765552 |  |  |  |  |  |
|  |  |  |  | *sichotense* Pojarkova | 100881 | **✓** | **✓** | **✓** |  | **✓** | **✓** | 2.47 | ZM | 1.316 | 2 | 1,289.680 |
|  |  | *Saluenensia* |  | *calostrotum* SPp. *keleticum* (Balfour & Forrest) Cullen | 100312 |  |  | **✓** |  |  |  |  |  |  |  |  |
|  |  |  |  | *saluenense* SPp. *saluenense* Balfour & Forrest | 100406 |  |  | **✓** |  |  |  |  |  |  |  |  |
|  |  | *Scabrifolia* |  | *racemosum* Franchet | OLD00795 | **✓** | **✓** | **✓** | **✓** |  |  | 3.92 | ZM | 1.474 | 2 | 1,444.520 |
|  |  |  |  | *spiciferum* | 100477 | **✓** | **✓** | **✓** | **✓** | **✓** | **✓** | 4.97 | ZM | 1.390 | 2 | 1,362.200 |
|  |  |  |  | *spinuliferum* Franchet | 100464 | **✓** | **✓** | **✓** | **✓** | **✓** | **✓** | 3.75 | ZM | 1.471 | 2 | 1,441.907 |
|  |  |  |  | *spinuliferum* Franchet | WTU 357197 |  |  |  | AY765651 | AY765652 | AY765654 |  |  |  |  |  |
|  |  | *Tephropepla* |  | *auritum* Tagg | 101052 (1293) | **✓** | **✓** | **✓** | **✓** | **✓** | **✓** | 4.28 | SP | 1.484 | 2 | 1,454.320 |
|  |  |  |  | *xanthostephanum* Merrill | 100474 | **✓** | **✓** | **✓** |  | **✓** |  | 4.07 | SP | 1.508 | 2 | 1,477.840 |
|  |  |  |  | *xanthostephanum* Merrill | WTU 357353 |  |  |  | AY765471 | AY765472 | AY765474 |  |  |  |  |  |
|  |  | *Trichoclada* |  | *mekongense* Franchet | WTU 357210 |  |  |  | AY765699 | AY765700 | AY765702 |  |  |  |  |  |
|  |  | *Triflora* |  | *ambiguum* a Hemsley | 100007 | **✓** | **✓** | **✓** | **✓** | **✓** | **✓** | 2.49 | ZM | 4.091 | 6 | 4,000.853 |
|  |  |  |  | *ambiguum* b Hemsley | OLD00801 | **✓** | **✓** | **✓** | **✓** | **✓** |  | 3.44 | ZM | 4.194 | 6 | 4,110.120 |
|  |  |  |  | *amesiae* Rehder & Wilson | 100009 |  |  | **✓** |  |  |  | 4.29 | ZM | 2.756 | 4 | 2,700.880 |
|  |  |  |  | *augustinii* SPp. *chasmanthum* Cullen | 100654 | **✓** | **✓** | **✓** |  |  |  | 2.16 | ZM | 3.764 | 4 | 3,688.720 |
|  |  |  |  | *concinnum* a Hemsley | 100326 | **✓** | **✓** | **✓** | **✓** | **✓** | **✓** | 2.89 | ZM | 4.239 | 6 | 4,154.547 |
|  |  |  |  | *concinnum* b Hemsley | 2008/375 | **✓** | **✓** | **✓** | **✓** | **✓** |  | 2.52 | ZM | 2.110 | 2 | 2,067.800 |
|  |  |  |  | *davidsonianum* Rehder & Wilson | 100329 | **✓** | **✓** | **✓** | **✓** | **✓** | **✓** | 2.88 | ZM | 3.721 | 4 | 3,646.907 |
|  |  |  |  | *keiskei* Miquel | 101358 |  |  | **✓** |  |  |  | 4.05 | ZM | 1.470 | 2 | 1,440.600 |
|  |  |  |  | *keiskei* Miquel | WTU 357242 |  |  |  | AY765687 | AY765688 | AY765690 |  |  |  |  |  |
|  |  |  |  | *lutescens* Franchet | 100362 | **✓** | **✓** | **✓** | **✓** | **✓** |  | 3.57 | ZM | 1.472 | 2 | 1,442.560 |
|  |  |  |  | *lutescens* Franchet | WTU 357245 |  |  |  | AY765675 | AY765676 | AY765678 |  |  |  |  |  |
|  |  |  |  | *pleistanthum* Balfour ex Wilding | 100471 | **✓** | **✓** | **✓** |  |  |  | 2.99 | ZM | 3.034 | 4 | 2,973.320 |
|  |  |  |  | *polylepis* Franchet | 100495 | **✓** | **✓** | **✓** |  |  |  | 4.03 | ZM | 1.498 | 2 | 1,468.040 |
|  |  |  |  | *rigidum* Franchet | 100498 | **✓** | **✓** | **✓** |  | **✓** |  | 2.26 | ZM | 2.992 | 4 | 2,932.160 |
|  |  |  |  | *siderophyllum* Franchet | WTU 357206 |  |  |  | AY765663 | AY765664 | AY765666 | 3.82 | ZM | 4.306 | 6 | 4,219.880 |
|  |  |  |  | *tatsienense* Franchet | 100424 |  |  | **✓** |  |  |  | 3.41 | ZM | 4.530 | 6 | 4,439.400 |
|  |  |  |  | *trichanthum* Rehder | 100803 | **✓** | **✓** | **✓** | **✓** | **✓** |  | 4.42 | ZM | 4.382 | 6 | 4,294.360 |
|  |  |  |  | *trichanthum* Rehder | WTU 357182 |  |  |  | AY765645 | AY765646 | AY765648 |  |  |  |  |  |
|  |  |  |  | *triflorum* Hooker | 1470 |  |  | **✓** |  |  |  | 2.87 | ZM | 4.264 | 6 | 4,178.720 |
|  |  |  |  | *triflorum* SPp. *triflorum* Hooker | WTU 357204 |  |  |  | AY765681 | AY765682 | AY765684 |  |  |  |  |  |
|  |  |  |  | *yunnanense* Franchet | 100449 | **✓** | **✓** | **✓** | **✓** | **✓** | **✓** | 3.20 | ZM | 4.363 | 4 | 4,276.067 |
|  |  |  |  | *zaleucum* var. *flaviflorum* Balfour & Smith | 2002/1096b | **✓** | **✓** | **✓** | **✓** | **✓** | **✓** | 5.30 | ZM | 1.091 | 2 | 1,068.853 |
|  |  | *Uniflora* |  | *pemakoense* Kingdon-Ward | 2008/389 |  |  | **✓** |  |  |  |  |  |  |  |  |
|  |  |  |  | *virgatum* Hooker | WTU 357220 |  |  |  | AY765669 | AY765670 | AY765672 |  |  |  |  |  |
|  | *Schistanthe* (*Vireya*) | *Discovireya* |  | *quadrasianum* var. *rosmarinifolium* | 1997/028 |  | **✓** | **✓** | **✓** |  | **✓** |  |  |  |  |  |
|  |  |  |  | *monodii* | 2007/609 |  |  | **✓** | **✓** |  | **✓** |  |  |  |  |  |
|  |  |  |  | *retusum* | 1973/064 | **✓** | **✓** | **✓** |  |  |  | 7.61 | HD | 1.418 | 2 | 1,389.640 |
|  |  | *Euvireya* | "*Albovireya*" | *aequabile* | 2002/372 |  | **✓** | **✓** | **✓** |  | **✓** |  |  |  |  |  |
|  |  |  |  | *aequabile* | RBGE |  | AY877284 |  |  |  |  |  |  |  |  |  |
|  |  |  |  | *aequabile* |  |  |  |  | GU445495 | GU445573 | GU445729 |  |  |  |  |  |
|  |  |  |  | *album* | 2005/1065 |  | **✓** | **✓** | **✓** |  |  | 6.81 | HD | 2.276 | 4 | 2,230.480 |
|  |  |  |  | *langulicarpum* | DB |  | AY877287 |  |  |  |  |  |  |  |  |  |
|  |  |  |  | *langulicarpum* |  |  |  |  | GU445500 | GU445578 | GU445734 |  |  |  |  |  |
|  |  |  |  | *zollingeri* | DB |  | AY877296 |  |  |  |  |  |  |  |  |  |
|  |  |  |  | *zollingeri* |  |  |  |  | GU445511 | GU445589 | GU445745 |  |  |  |  |  |
|  |  |  | "*Euvireya*" | *acrophilum* | 2002/371 |  | **✓** | **✓** |  |  |  |  |  |  |  |  |
|  |  |  |  | *aurigeranum* | 1986/003 | **✓** | **✓** | S2 |  |  |  | 3.23 | HD | 3.312 | 4 | 3,245.760 |
|  |  |  |  | *blackii* | 2002/373 |  | **✓** | **✓** | **✓** |  |  |  |  |  |  |  |
|  |  |  |  | *burtii* | 2000/125 |  | **✓** | **✓** | **✓** |  |  | 7.49 | ZM | 2.822 | 4 | 2,765.560 |
|  |  |  |  | *celebicum* | 2002/374 |  | **✓** | **✓** | **✓** |  |  |  |  |  |  |  |
|  |  |  |  | *christi* | LC |  | AY877269 |  |  |  |  |  |  |  |  |  |
|  |  |  |  | *christi* |  |  |  |  | GU445467 | GU445545 | GU445701 |  |  |  |  |  |
|  |  |  |  | *christi* | 1997/015 |  | **✓** | S2 |  |  |  |  |  |  |  |  |
|  |  |  |  | *christianae* | 2002/375 | **✓** | **✓** | S2 |  |  |  |  |  |  |  |  |
|  |  |  |  | *citrinum* var. *citrinum* | 2005/1211 |  | **✓** | **✓** |  |  |  |  |  |  |  |  |
|  |  |  |  | *craSPifolium* | 1997/051 |  | **✓** | **✓** | **✓** | **✓** |  |  |  |  |  |  |
|  |  |  |  | *craSPifolium* Stapf | RSF 73 |  |  |  | AY765603 | AY765604 | AY765606 |  |  |  |  |  |
|  |  |  |  | *culminicola* var. *culminicola* | 1996/011 |  | **✓** | **✓** | **✓** |  |  |  |  |  |  |  |
|  |  |  |  | *gracilentum* | 1997/016 |  | **✓** |  |  |  |  | 4.11 | ZM | 2.742 | 4 | 2,687.160 |
|  |  |  |  | *gracilentum* | LC |  | AY877271 |  |  |  |  |  |  |  |  |  |
|  |  |  |  | *gracilentum* |  |  |  |  | GU445471 | GU445549 | GU445705 |  |  |  |  |  |
|  |  |  |  | *javanicum* Bennett | NA |  |  | AB012742 |  |  |  |  |  |  |  |  |
|  |  |  |  | *javanicum* SPp. *javanicum* | 1997/021 |  | **✓** | **✓** | **✓** | **✓** |  | 3.21 | HD | 2.036 | 4 | 1,995.280 |
|  |  |  |  | *laetum* | 1973/020 | **✓** | **✓** | **✓** |  |  |  | 2.91 | SP | 3.400 | 4 | 3,332.000 |
|  |  |  |  | *leucogigas* | 2000/147 |  | **✓** | **✓** | **✓** | **✓** |  |  |  |  |  |  |
|  |  |  |  | *lochiae* | LC |  | AY877279 |  |  |  |  |  |  |  |  |  |
|  |  |  |  | *lochiae* |  |  |  |  | GU445460 | GU445538 | GU445694 |  |  |  |  |  |
|  |  |  |  | *lochiae*/*viriolosum* | 1972/109 | **✓** | **✓** | S2 |  |  |  | 5.01 | HD | 2.086 | 4 | 2,044.280 |
|  |  |  |  | *luraluense* | LC |  | AY877277 |  |  |  |  |  |  |  |  |  |
|  |  |  |  | *luraluense* |  |  |  |  | GU445646 | GU445542 | GU445698 |  |  |  |  |  |
|  |  |  |  | *macgregoriae* | 2007/614 | **✓** | **✓** | **✓** |  |  |  | 3.36 | HD | 3.200 | 4 | 3,136.000 |
|  |  |  |  | *multicolor* | 1997/026 |  | **✓** | **✓** | **✓** | **✓** |  |  |  |  |  |  |
|  |  |  |  | *nieuwenhuisii* | 2005/1208 |  | **✓** | **✓** | **✓** |  |  |  |  |  |  |  |
|  |  |  |  | *orbiculatum* | 2002/386 |  | **✓** | **✓** |  |  |  | 3.80 | HD | 2.190 | 4 | 2,146.200 |
|  |  |  |  | *polyanthemum* | 1994/013 | **✓** | **✓** | **✓** | **✓** | **✓** | **✓** | 2.80 | SP | 2.232 | 4 | 2,187.360 |
|  |  |  |  | *praetervisum* | 1997/057 |  | **✓** | **✓** | **✓** |  | **✓** |  |  |  |  |  |
|  |  |  |  | *renschieanum* | 2002/049 |  | **✓** | **✓** | **✓** | **✓** |  |  |  |  |  |  |
|  |  |  |  | *rhodopus* | BCJR129 |  | AY877293 |  |  |  |  |  |  |  |  |  |
|  |  |  |  | *rhodopus* |  |  |  |  | GU445509 | GU445587 | GU445743 |  |  |  |  |  |
|  |  |  |  | *rousei* | DB |  | AY877291 |  |  |  |  |  |  |  |  |  |
|  |  |  |  | *rousei* |  |  |  |  | GU445492 | GU445570 | GU445726 |  |  |  |  |  |
|  |  |  |  | *rousei* | 2002/362 |  | **✓** | **✓** |  |  |  |  |  |  |  |  |
|  |  |  |  | *rubineiflorum* | LC |  | AY877294 |  |  |  |  |  |  |  |  |  |
|  |  |  |  | *rubineiflorum* |  |  |  |  | GU445473 | GU445551 | GU445707 |  |  |  |  |  |
|  |  |  |  | *rugosum* | 2002/391 |  |  | **✓** |  |  |  |  |  |  |  |  |
|  |  |  |  | *saxifragoides* | DB |  | AY877299 |  |  |  |  |  |  |  |  |  |
|  |  |  |  | *stenophyllum* var. *angustifolium* | 101800 |  | **✓** | **✓** | **✓** |  | **✓** |  |  |  |  |  |
|  |  |  |  | *taxifolium* | 2000/144 |  | **✓** | **✓** | **✓** |  |  | 4.36 | HD | 2.314 | 4 | 2,267.720 |
|  |  |  |  | *vanvuurenii* | 2007/627 |  |  | **✓** |  |  |  |  |  |  |  |  |
|  |  |  |  | *verticillatum* | 2005/1210 |  | **✓** | **✓** |  |  |  | 5.16 | HD | 2.120 | 4 | 2,077.600 |
|  |  |  |  | *viriosum* | LC |  | AY877288 |  |  |  |  |  |  |  |  |  |
|  |  |  |  | *wrightianum* var. *wrightianum* | 2006/376 |  | **✓** | **✓** | **✓** |  | **✓** |  |  |  |  |  |
|  |  |  |  | *zoelleri* | LC |  | AY877302 |  |  |  |  |  |  |  |  |  |
|  |  |  |  | *zoelleri* |  |  |  |  | GU445469 | GU445547 | GU445703 |  |  |  |  |  |
|  |  |  | "*Phaeovireya*" | *caliginis* | 1997/014 |  | **✓** | S2 |  |  |  |  |  |  |  |  |
|  |  |  |  | *dianthosmum* | 2002/378 |  | **✓** | S2 |  |  |  |  |  |  |  |  |
|  |  |  |  | *dielsianum* Schlechter | RSF 95 |  |  |  | AY765579 | AY765580 | AY765582 |  |  |  |  |  |
|  |  |  |  | *dielsianum* var. *dielsianum* | 2002/379 |  | **✓** | S2 |  |  |  |  |  |  |  |  |
|  |  |  |  | *gardenia* | 2006/549 |  | **✓** | **✓** | **✓** |  |  |  |  |  |  |  |
|  |  |  |  | *hellwigii* | 2007/610 |  | **✓** | S2 |  |  |  |  |  |  |  |  |
|  |  |  |  | *konori* | 2001/003 | **✓** | **✓** | **✓** |  |  |  | 11.36 | HD | 3.062 | 4 | 3,000.760 |
|  |  |  |  | *konori* Beccari | WTU 357257 |  |  |  | AY765597 | AY765598 | AY765600 |  |  |  |  |  |
|  |  |  |  | *leptanthum* | 2000/132 |  | **✓** | **✓** | **✓** | **✓** |  |  |  |  |  |  |
|  |  |  |  | *leptanthum* | LC |  | AY877275 |  |  |  |  |  |  |  |  |  |
|  |  |  |  | *phaeochitum* | 1999/155 |  | **✓** | S2 |  |  |  | 4.94 | SP | 3.542 | 4 | 3,471.160 |
|  |  |  |  | *rarum* | LC |  | AY877285 |  |  |  |  |  |  |  |  |  |
|  |  |  |  | *rarum* | X/20125 |  |  | **✓** |  |  |  |  |  |  |  |  |
|  |  |  |  | *ericoides* | LC |  | AY877270 |  |  |  |  |  |  |  |  |  |
|  |  |  |  | *ericoides* |  |  |  |  | GU445526 | GU445604 | GU445760 |  |  |  |  |  |
|  |  |  | "*Siphonovireya*" | *agathodaemonis* | 2008/189 |  | **✓** | **✓** | **✓** |  | **✓** |  |  |  |  |  |
|  |  |  |  | *herzogii* Warburg | 2002/382 |  | **✓** | S2 |  |  |  |  |  |  |  |  |
|  |  |  |  | *herzogii Warburg* | LC |  | AY877272 |  |  |  |  |  |  |  |  |  |
|  |  |  |  | *herzogii* Warburg | WTU 357185 |  |  |  | AY765591 | AY765592 | AY765594 |  |  |  |  |  |
|  |  |  | "*Solenovireya*" | *alborugosum* | 2005/1063 |  | **✓** | **✓** | **✓** |  | **✓** |  |  |  |  |  |
|  |  |  |  | *alborugosum* | DB |  | AY877283 |  |  |  |  |  |  |  |  |  |
|  |  |  |  | *alborugosum* |  |  |  |  | GU445480 | GU445558 | GU445714 |  |  |  |  |  |
|  |  |  |  | *armitii* | 2000/091 |  | **✓** | **✓** |  |  |  |  |  |  |  |  |
|  |  |  |  | *carringtoniae* | 1999/154 |  | **✓** | **✓** | **✓** |  |  |  |  |  |  |  |
|  |  |  |  | *cruttwellii* | 2002/398 |  | **✓** |  |  |  |  | 5.63 | HD | 2.778 | 4 | 2,722.440 |
|  |  |  |  | *edanoi* |  |  |  |  | GU445485 | GU445563 | GU445719 |  |  |  |  |  |
|  |  |  |  | *edanoi SPp. pneumonanthum* | DB |  | AY877282 |  |  |  |  |  |  |  |  |  |
|  |  |  |  | *edanoi* SPp. *pneumonanthum* | 2002/388 | **✓** | **✓** | **✓** |  |  |  |  |  |  |  |  |
|  |  |  |  | *jasminiflorum* | 2000/130 |  | **✓** | **✓** | **✓** | **✓** |  |  |  |  |  |  |
|  |  |  |  | *jasminiflorum* |  |  |  |  | GU445481 | GU445559 | GU445715 |  |  |  |  |  |
|  |  |  |  | *jasminiflorum* var. *heuSPeri* | LC |  | AY877273 |  |  |  |  |  |  |  |  |  |
|  |  |  |  | *loranthiflorum* | 1997/025 | **✓** | **✓** | S2 |  |  |  |  |  |  |  |  |
|  |  |  |  | *loranthiflorum* | LC |  | AY877276 |  |  |  |  |  |  |  |  |  |
|  |  |  |  | *loranthiflorum* |  |  |  |  | GU445478 | GU445556 | GU445712 |  |  |  |  |  |
|  |  |  |  | *majus* | 2003/036 |  | **✓** | **✓** | **✓** | **✓** |  |  |  |  |  |  |
|  |  |  |  | *radians* Smith | WTU 357163 |  |  |  | AY765585 | AY765586 | AY765588 |  |  |  |  |  |
|  |  |  |  | *ruttenii* | LC |  | AY877295 |  |  |  |  |  |  |  |  |  |
|  |  |  |  | *ruttenii* |  |  |  |  | GU445483 | GU445561 | GU445717 |  |  |  |  |  |
|  |  |  |  | *tuba* | 2002/040 |  | **✓** | **✓** |  |  | **✓** |  |  |  |  |  |
|  |  |  |  | *tuba* | LC |  | AY877300 |  |  |  |  |  |  |  |  |  |
|  |  |  |  | *tuba* |  |  |  |  | GU445459 | GU445537 | GU445693 |  |  |  |  |  |
|  |  | *Malayovireya* |  | *apoanum* | 1999/073 |  | **✓** | **✓** |  |  |  | 5.06 | HD | 2.818 | 4 | 2,761.640 |
|  |  |  |  | *apoanum* | RBGE |  | AY877267 |  |  |  |  | 7.76 | ZM | 2.850 | 4 | 2,793.000 |
|  |  |  |  | *apoanum* |  |  |  |  | GU445523 | GU445601 | GU445757 |  |  |  |  |  |
|  |  |  |  | *fallacinum* | 2005/1205 |  | **✓** | **✓** | **✓** | **✓** |  |  |  |  |  |  |
|  |  |  |  | *himantodes* var. *himantodes* | 1997/020 | **✓** | **✓** | **✓** |  | **✓** |  |  |  |  |  |  |
|  |  |  |  | *malayanum* | 2007/613 |  | **✓** | **✓** | **✓** | **✓** |  |  |  |  |  |  |
|  |  |  |  | *malayanum* | DB |  | AY877278 |  |  |  |  |  |  |  |  |  |
|  |  |  |  | *malayanum* |  |  |  |  | GU445521 | GU445599 | GU445755 |  |  |  |  |  |
|  |  | *Pseudovireya* |  | *asperulum* Hutchinson & Kingdon-Ward | WTU 357187 |  |  |  | AY765615 | AY765616 | AY765618 |  |  |  |  |  |
|  |  |  |  | *emarginatum* | 1997/094 | **✓** | **✓** | **✓** | **✓** | **✓** | **✓** | 7.55 | ZM | 2.774 | 4 | 2,718.520 |
|  |  |  |  | *kawakamii* Hayata | X/20117 |  | **✓** | **✓** |  |  |  | 2.88 | HD | 1.372 | 2 | 1,344.560 |
|  |  |  |  | *kawakamii* Hayata | RSF 79-026 |  | GU176635 |  |  |  |  |  |  |  |  |  |
|  |  |  |  | *kawakamii* Hayata |  |  |  |  | GU445353 | GU445613 | GU445769 |  |  |  |  |  |
|  |  |  |  | *nanophyton* | BCJ 46 |  | AY877290 |  |  |  |  |  |  |  |  |  |
|  |  |  |  | *nanophyton* |  |  |  |  | GU445524 | GU445602 | GU445758 |  |  |  |  |  |
|  |  |  |  | *quadrasianum* var. *rosmariniflorum* | DB |  | AY877292 |  |  |  |  |  |  |  |  |  |
|  |  |  |  | *retusum* | LC |  | AY877286 |  |  |  |  |  |  |  |  |  |
|  |  |  |  | *retusum* |  |  |  |  | GU445530 | GU445608 | GU445764 |  |  |  |  |  |
|  |  |  |  | *rushfortii* | 2002/392 |  | **✓** | **✓** | **✓** | **✓** |  | 5.54 | HD | 1.354 | 2 | 1,326.920 |
|  |  |  |  | *santapaui* | OLD00886 |  | **✓** | **✓** |  | **✓** |  |  |  |  |  |  |
|  |  |  |  | *santapaui* Sastry et al. | NA |  |  | AB012743 |  |  |  |  |  |  |  |  |
|  |  |  |  | *santapaui* Sastry et al. | WTU 357211 |  |  |  | AY765621 | AY765622 | AY765624 |  |  |  |  |  |
|  |  |  |  | *sororium* | 1994/097 |  | **✓** | **✓** | **✓** | **✓** | **✓** |  |  |  |  |  |
|  |  |  |  | *sororium* Sleumer | WTU 357221 |  |  |  | AY765609 | AY765610 | AY765612 |  |  |  |  |  |
|  |  |  |  | *vaccinioides* Hooker | WTU 357212 |  |  |  | AY765627 | AY765628 | AY765630 |  |  |  |  |  |
| *Therorhodion* |  |  |  | *camtschaticum* a Pallas | RSF 73-054 |  | GU176637 |  |  |  |  |  |  |  |  |  |
|  |  |  |  | *camtschaticum* Pallas | NA |  |  | AB012744 |  |  |  |  |  |  |  |  |
|  |  |  |  | *camtschaticum* Pallas | WTU 357178 |  |  |  | AY765981 | AY765982 | AY765984 |  |  |  |  |  |
|  |  |  |  | *camtschaticum* SPp. *camtschaticum* Pallas | 100317 | **✓** | **✓** | **✓** |  |  |  | 5.73 | ZM | 1.165 | 2 | 1,141.373 |
| *Tsutsusi* | *Brachycalyx* |  |  | *dilatatum* Miquel | 100904 | **✓** | **✓** | **✓** | **✓** | **✓** | **✓** | 4.99 | ZM | 1.234 | 2 | 1,209.320 |
|  |  |  |  | *dilatatum* Miquel | E 1975-0766B **?** |  | EU855852 |  |  |  |  |  |  |  |  |  |
|  |  |  |  | *farrerae* Tate ex Sweet | RSF 78-037 |  | EU855854 |  |  |  |  |  |  |  |  |  |
|  |  |  |  | *farrerae* Tate ex Sweet | NA |  |  | AB012745 |  |  |  |  |  |  |  |  |
|  |  |  |  | *kiyosumense* (Makino) Makino | RSF 77-027 |  | EU855858 |  |  |  |  |  |  |  |  |  |
|  |  |  |  | *lagopus* Nakai var. lagopus | RSF 03-432 |  | EU855859 |  |  |  |  |  |  |  |  |  |
|  |  |  |  | *mariesii* Hemsley & Wilson | WTU 357222 |  |  |  | AY765903 | AY765904 | AY765906 |  |  |  |  |  |
|  |  |  |  | *mayebarae* Nakai & Hara | E 1995-0441A |  | EU855860 |  |  |  |  |  |  |  |  |  |
|  |  |  |  | *reticulatum* Don ex Don | 100400 |  | **✓** | **✓** |  |  | **✓** |  |  |  |  |  |
|  |  |  |  | *reticulatum* Don ex Don | E 1975-2245 |  | EU855864 |  |  |  |  |  |  |  |  |  |
|  |  |  |  | *sanctum* Nakai | RSF 73-250 |  | EU855866 |  |  |  |  |  |  |  |  |  |
|  |  |  |  | *viscistylum* Nakai | RSF 77-028 |  | EU855876 |  |  |  |  |  |  |  |  |  |
|  |  |  |  | *wadanum* Makino | 100436 | **✓** | **✓** | **✓** | **✓** | **✓** | **✓** |  |  |  |  |  |
|  |  |  |  | *wadanum* a Makino | E 1976-1072D |  | EU855877 |  |  |  |  |  |  |  |  |  |
|  |  |  |  | *wadanum* Makino | NA |  |  | AB012746 |  |  |  |  |  |  |  |  |
|  |  |  |  | *wadanum* Makino | WTU 357190 |  |  |  | AY765897 | AY765898 | AY765900 |  |  |  |  |  |
|  |  |  |  | *wadanum* Makino |  | AF452218 |  |  |  |  |  |  |  |  |  |  |
|  |  |  |  | *weyrichii* Maximowicz | E 1994-2387A |  | EU855878 |  |  |  |  |  |  |  |  |  |
|  |  |  |  | *breviperulatum* Hayata | RSF 82-088 |  | EU855851 |  |  |  |  |  |  |  |  |  |
|  | *Tsutsusi* |  |  | *eriocarpum* (Hayata) Nakai | 100650 | **✓** | **✓** | **✓** | **✓** | **✓** |  | 3.49 | ZM | 1.509 | 2 | 1,478.493 |
|  |  |  |  | *eriocarpum* (Hayata) Nakai | E 1988-0983 |  | EU855853 |  |  |  |  |  |  |  |  |  |
|  |  |  |  | *indicum* (Linneaus) Sweet | E 1995-1015A |  | EU855855 |  |  |  |  |  |  |  |  |  |
|  |  |  |  | *indicum* (Linneaus) Sweet | NA |  |  | AB012747 |  |  |  |  |  |  |  |  |
|  |  |  |  | *kaempferi* Planchon | 100357 | **✓** | **✓** | **✓** | **✓** | **✓** |  | 5.69 | ZM | 1.142 | 2 | 1,119.160 |
|  |  |  |  | *kiusianum* Makino | RSF 79-059 - RBGE 191029 |  | EU855857 |  |  |  |  |  |  |  |  |  |
|  |  |  |  | *mucronatum* (Blume) G. Don |  |  | AF393412 |  |  |  |  |  |  |  |  |  |
|  |  |  |  | *mucronatum* (Blume) G. Don var. *ripense* (Makino) Wilson | RSF 98-244 |  | EU855861 |  |  |  |  |  |  |  |  |  |
|  |  |  |  | *mucronatum* (Blume) G. Don var. *ripense* (Makino) Wilson | 100403 | **✓** | **✓** | **✓** |  |  |  | 3.62 | ZM | 1.552 | 2 | 1,520.960 |
|  |  |  |  | *nakaharae* Hayata | RSF 74-85 |  | EU855862 |  |  |  |  |  |  |  |  |  |
|  |  |  |  | *obtusum* var. *kaempferi* | E 1976-1898 |  | EU855856 |  |  |  |  |  |  |  |  |  |
|  |  |  |  | *obtusum* var. *kaempferi* | NA |  |  | AB012748 |  |  |  |  |  |  |  |  |
|  |  |  |  | *oldhamii* Maximowicz | 100871 |  | **✓** |  |  |  |  | 5.57 | ZM | 1.254 | 2 | 1,228.920 |
|  |  |  |  | *oldhamii* Maximowicz | E 1971-0104 |  | EU855863 |  |  |  |  |  |  |  |  |  |
|  |  |  |  | *rubropilosum* Hayata | RSF 73-242 |  | EU855865 |  |  |  |  |  |  |  |  |  |
|  |  |  |  | *rubropilosum* Hayata | WTU 357223 |  |  |  | AY765909 | AY765910 | AY765912 |  |  |  |  |  |
|  |  |  |  | *scabrum* Don | RSF 87-062 |  | EU855867 |  |  |  |  |  |  |  |  |  |
|  |  |  |  | *serpyllifolium* (Gray) Miquel | RSF 76-356 |  | EU855868 |  |  |  |  |  |  |  |  |  |
|  |  |  |  | *simsii* Planchon | NA |  | EU855869 |  |  |  |  |  |  |  |  |  |
|  |  |  |  | *stenopetalum* (Hogg) Mabberley | RSF 37-34 |  | EU855870 |  |  |  |  |  |  |  |  |  |
|  |  |  |  | *subseSPile* Rendle | RSF 99-310 |  | EU855871 |  |  |  |  |  |  |  |  |  |
|  |  |  |  | *tashiroi* Maximowicz | 100422 | **✓** | **✓** | **✓** | **✓** | **✓** | **✓** | 4.80 | ZM | 1.222 | 2 | 1,197.560 |
|  |  |  |  | *tashiroi* Maximowicz | RSF - s.n. |  | EU855872 |  |  |  |  |  |  |  |  |  |
|  |  |  |  | *tashiroi* Maximowicz | NA |  |  | AB012749 |  |  |  |  |  |  |  |  |
|  |  |  |  | *tashiroi* Maximowicz | WTU 357213 |  |  |  | AY765891 | AY765892 | AY765894 |  |  |  |  |  |
|  |  |  |  | *tosaense* Makino | RSF 98-585 |  | EU855873 |  |  |  |  |  |  |  |  |  |
|  |  |  |  | *tschonoskii* Maximowicz | E 1975-0766B ? |  | EU855874 |  |  |  |  |  |  |  |  |  |
|  |  |  |  | *tsusiophyllum* Sugimoto | K 1985-4676 |  | GU176636 |  |  |  |  |  |  |  |  |  |
|  |  |  |  | *tsutsuiphyllum* Sugimoto | NA |  |  | AB012750 |  |  |  |  |  |  |  |  |
|  |  |  |  | *tsutsuiphyllum* Sugimoto | WTU 357177 |  |  |  | AY765927 | AY765928 | AY765930 |  |  |  |  |  |
|  |  |  |  | *yedoense* Maximowicz var. *poukhanense* (Léveillé) Nakai | E 1977-2159B |  | EU855879 |  |  |  |  |  |  |  |  |  |
| **Outgroup** |  |  |  |  |  |  |  |  |  |  |  |  |  |  |  |  |
|  |  | *Bejaria* |  | *racemosa* |  |  | BRU48604 |  |  |  |  |  |  |  |  |  |
|  |  | *Calluna* |  | *vulgaris* (Linneaus) Hull | OLD06730 | **✓** | **✓** | **✓** |  | **✓** |  |  |  |  |  |  |
|  |  | *CaSPiope* |  | *lycopodioides* |  |  |  | AB012754 |  |  |  |  |  |  |  |  |
|  |  | *Elliottia* |  | *paniculata* |  |  |  | AB012753 |  |  |  |  |  |  |  |  |
|  |  | *Elliottia* |  | *racemosa* |  |  | ERU48582 |  |  |  |  |  |  |  |  |  |
|  |  | *Empetrum* |  | *nigrum* Linneaus | OLD06731 | **✓** | **✓** |  |  | **✓** |  |  |  |  |  |  |
|  |  | *Empetrum* |  | *nigrum* Linneaus |  |  | GU176626 |  |  |  |  |  |  |  |  |  |
|  |  | *Kalmia* |  | *angustifolia* Linneaus | OLD06732 | **✓** | **✓** | **✓** |  | **✓** |  |  |  |  |  |  |
|  |  | *Kalmia* |  | *angustifolia* Linneaus |  |  | KAU48599 |  |  |  |  |  |  |  |  |  |
|  |  | *Kalmia* |  | *procumbens* (Linneaus) Desvaux = *Kalmia procumbens* Desvaux | OLD06733 | **✓** | **✓** |  |  |  |  |  |  |  |  |  |
|  |  | *Kalmia* |  | *procumbens* (Linneaus) Desvaux = *Kalmia procumbens* Desvaux |  |  | LPU48610 |  |  |  |  |  |  |  |  |  |
|  |  | *Phyllodoce* |  | *empetriformis* (Smith) Don | OLD00930 | **✓** | **✓** | **✓** |  |  |  |  |  |  |  |  |
|  |  | *Phyllodoce* |  | *nipponica* |  |  | PNU48606 |  |  |  |  |  |  |  |  |  |
|  |  | *Vaccinium* |  | x *intermedium* Ruthe | OLD06734 | **✓** | **✓** | S2 |  |  |  |  |  |  |  |  |
|  |  | *Vaccinium* |  | *myrtilloides* Michaux | OLD06735 | **✓** |  |  |  |  |  |  |  |  |  |  |
