## Supplemental Table 3 for "Incongruent phylogenies and its implications for the study of diversification, taxonomy and genome size evolution of *Rhododendron* (Ericaceae)"

| **TABLE S3.** PCR reaction profiles for the regions investigated; function of the different thermal reaction steps | | | | |
| --- | --- | --- | --- | --- |
| **Region** | **Temperatur** | **Time** | **Cycles** | **Function** |
| ITS | 94 °C | 1 min | 1 | Initial Denaturation |
|  | 94 °C | 18 sec |  | Denaturation |
|  | 55 °C | 30 sec | 35 | Annealing |
|  | 72 °C | 1 min |  | Extension |
|  | 72 °C | 8 min | 1 | Final Extension |
| *trn*LF | 94 °C | 1 min | 1 | Initial Denaturation |
|  | 94 °C | 30 sec |  | Denaturation |
|  | 54 °C | 30 sec | 35 | Annealing |
|  | 72 °C | 1 min 20 sec |  | Extension |
|  | 72 °C | 10 min | 1 | Final Extension |
| *trn*K-*mat*K | 94 °C | 2 min | 1 | Initial Denaturation |
|  | 94 °C | 30 sec |  | Denaturation |
|  | 55 °C | 1 min | 30 | Annealing |
|  | 72 °C | 1 min |  | Extension |
|  | 72 °C | 7 min |  | Final Extension |
| *rpb*2-*i* | 94 °C | 4 min | 1 | Initial Denaturation |
|  | 94 °C | 45 sec |  | Denaturation |
|  | 58.80 °C | 45 sec | 35 | Annealing |
|  | 72 °C | 1 min 20 sec |  | Extension |
|  | 72 °C | 10 min | 1 | Final Extension |
